# Safety and biodistribution of Nanoligomers^™^ targeting SARS-CoV-2 genome for treatment of COVID-19

**DOI:** 10.1101/2022.07.19.500688

**Authors:** Colleen R. McCollum, Colleen M. Courtney, Nolan J. O’Connor, Thomas R. Aunins, Tristan X. Jordan, Keegan Rogers, Stephen Brindley, Jared M. Brown, Prashant Nagpal, Anushree Chatterjee

**Author notes:** Corresponding author: Anushree Chatterjee.

## Abstract

As the world braces to enter its third year in the coronavirus disease 2019 (COVID-19) pandemic, the need for accessible and effective antiviral therapeutics continues to be felt globally. The recent surge of Omicron variant cases has demonstrated that vaccination and prevention alone cannot quell the spread of highly transmissible variants. A safe and nontoxic therapeutic with an adaptable design to respond to the emergence of new variants is critical for transitioning to treatment of COVID-19 as an endemic disease. Here, we present a novel compound, called SBCoV202, that specifically and tightly binds the translation initiation site of RNA-dependent RNA polymerase within the severe acute respiratory syndrome coronavirus 2 (SARS-CoV-2) genome, inhibiting viral replication. SBCoV202 is a Nanoligomer,^™^ a molecule that includes peptide nucleic acid sequences capable of binding viral RNA with single-base-pair specificity to accurately target the viral genome. The compound has been shown to be safe and nontoxic in mice, with favorable biodistribution, and has shown efficacy against SARS-CoV-2 *in vitro*. Safety and biodistribution were assessed after three separate administration methods, namely intranasal, intravenous, and intraperitoneal. Safety studies showed the Nanoligomer caused no outward distress, immunogenicity, or organ tissue damage, measured through observation of behavior and body weight, serum levels of cytokines, and histopathology of fixed tissue, respectively. SBCoV202 was evenly biodistributed throughout the body, with most tissues measuring Nanoligomer concentrations well above the compound K_D_ of 3.37 nM. In addition to favorable availability to organs such as the lungs, lymph nodes, liver, and spleen, the compound circulated through the blood and was rapidly cleared through the renal and urinary systems. The favorable biodistribution and lack of immunogenicity and toxicity set Nanoligomers apart from other antisense therapies, while the adaptability of the nucleic acid sequence of Nanoligomers provides a defense against future emergence of drug resistance, making these molecules an attractive potential treatment for COVID-19.

## Background

Early in 2020, severe acute respiratory syndrome coronavirus 2 (SARS-CoV-2) began a rapid spread across the world, highlighting the systemic inadequacies to current antiviral options.^1^ While vaccines were rapidly developed and distributed,^2–4^ the transmission of the virus remains high in unvaccinated populations. Emerging variants of concern such as Delta and Omicron introduce new challenges to the control of spread and treatment of infected individuals. Further, the efficacy of currently available vaccines against emerging variants is not guaranteed, with signs that individuals may still be susceptible to infection by the Omicron variant with a full two-dose vaccination.^5^ As nations around the globe struggled to prevent the transmission of the disease and quell the rising fatalities, it became starkly apparent that effective and well-researched antivirals for coronaviruses were greatly limited. Monoclonal antibody treatments have been approved on an emergency basis by the Food and Drug Administration,^6^ but the optimal timing of administration remains at the difficult-to-detect stage of early and non-severe disease progression, and the risk of resistance emergence poses a concern for widespread use.^7–9^ Remdesivir, an antiviral developed for Ebola and Marburg viral infections, targets the RNA-dependent RNA polymerase and is able to stall its action in SARS-CoV-2.^10,11^ While clinical outcomes of patients treated with remdesivir were promising,^12,13^ side effects of remdesivir treatment can be severe.^14^ Even with use of these treatments, 13.6% of COVID-19 cases that require hospitalization are fatal in the U.S.^15^ Further, the risk remains of the emergence of a variant that is resistant to current treatment options. There is thus an urgent need for new, adaptable treatments for SARS-CoV-2 infection. Here, we present a compound based on antisense technology that can specifically and effectively target SARS-CoV-2 and prevent viral replication. Use of this compound could prevent severe disease caused by SARS-CoV-2, and the compound can be easily adapted to target emerging variant strains.

Infection of the body by SARS-CoV-2 causes coronavirus disease 2019 (COVID-19) in humans. Infection typically begins in the upper respiratory tract, and spreads to the lungs. Symptoms of COVID-19 include cough, fever, and labored breathing, but the virus may also trigger the host immune system to overreact, resulting in cytokine storm syndrome (CSS). The combination of damage from the virus and CSS can lead to acute respiratory distress syndrome, uncontrolled inflammation, and organ failure.^16^ To date, over 437 million infections and nearly 6 million deaths have occurred due to COVID-19 globally. While vaccines have significantly slowed the spread of SARS-CoV-2, they cannot prevent disease with 100% efficacy, especially when vaccine rollouts are delayed or subsets of the population otherwise elect out of vaccination. Preventing viral replication would slow both the spread of the virus through populations and limit cases of severe infections.

SARS-CoV-2 is a coronavirus with a positive single-stranded RNA genome.^16,17^ Viral particles are enveloped, and spike proteins on their surfaces, which are recognized by human angiotensin-converting enzyme 2 (hACE2) receptors, assisted by transmembrane protease receptor, serine 2 (TMPRSS2),^18^ allow for entry into host cells and subsequent infection. Once within a mammalian host cell, the viral genome is released. Host cell machinery is used to translate viral proteins, including RNA-dependent RNA polymerase (RdRp), one of the main enzymes responsible for the replication of the viral genome. The RNA polymerase then transcribes copies of the genome to populate newly assembled viral particles, which are released from infected cells to spread infection.^16,17^

Currently available antivirals for COVID-19 largely do not act upon the viral genome. Some antivirals being used clinically bind the RdRp, including remdesivir.^11,12^ These antivirals have been applied following emergency approval by the FDA, though cases of severe side effects have hampered their success and the possibility of emergence of resistant variants remains a threat.^14^ Monoclonal antibody therapy boosts the body’s antibody response to clear infection, though the infection must be detected early for significant benefit.^19^ While these treatment options remain the best options for physicians currently, a more robust selection of treatments would bolster the global response to the pandemic.

A variety of approaches have been utilized to target the SARS-CoV-2 genome and prevent viral replication. Promising studies using clustered regularly interspaced short palindromic repeats (CRISPR) have targeted conserved sequences across a range of coronaviruses.^20,21^ The translation of this technology awaits development of an effective delivery mechanism. Short interfering RNA (siRNA) have similarly been designed to target conserved sequences in SARS-CoV-2 and also face challenges of delivery, which will likely be achieved through the use of lipid nanoparticles.^22^ Lipid nanoparticles, however, have been shown to cause innate immune response,^23,24^ and further optimization will be necessary to ensure that the hyper-responsive immune systems of patients suffering from CSS are not negatively affected by this delivery method. Other antisense oligonucleotides designed to directly target the SARS-CoV-2 genome bind untranslated regions or transcriptional regulatory sequences and are early in development.^25^ Direct-acting antivirals (DAAs) such as these are beneficial as treatment options as they do not rely on the activity of the host immune system and can act immediately upon any present viral particles rather than depending on administration at optimal infection progression.^26^

The antiviral compound presented here, called SBCoV202, was developed by Sachi Bioworks, Inc, and relies on antisense technology. In Nanoligomer^™^ molecules such as SBCoV202, a short peptide nucleic acid (PNA) sequence, conjugated to a gold nanoparticle, can specifically and tightly bind a target sequence of RNA (Figure 1A). PNAs are oligomers comprised of amine-nucleotide subunits. This creates a nucleic acid sequence with a rigid, stable amine backbone and a nucleotide sequence capable of highly specific binding to a target.^27^ The PNAs used in these Nanoligomers were rationally designed to target the translation initiation site of the RNA-dependent RNA polymerase within the SARS-CoV-2 genome while minimizing off-target binding within the host genome. A PNA molecule developed by Li et al., designed to target the translation initiation site with the use of a cell-penetrating peptide (CPP) to aid transport, showed efficacy in reducing SARS-CoV-2 infection.^28^ The addition of the gold nanoparticle improves transport of the PNA molecule into cells. Innate immune response has been observed with other methods to increase transport of antisense therapies, including PNA conjugated to CPP,^29,30^ siRNA,^31,32^ liposomes, and lipid-encapsulation.^24,33^ Through the use of biocompatible gold nanoparticles, Nanoligomers circumvent concerns faced by other antisense therapies and their delivery mechanisms by inducing no innate immune response and being easily transported across membranes without dependence on endocytosis.

**Figure 1:**
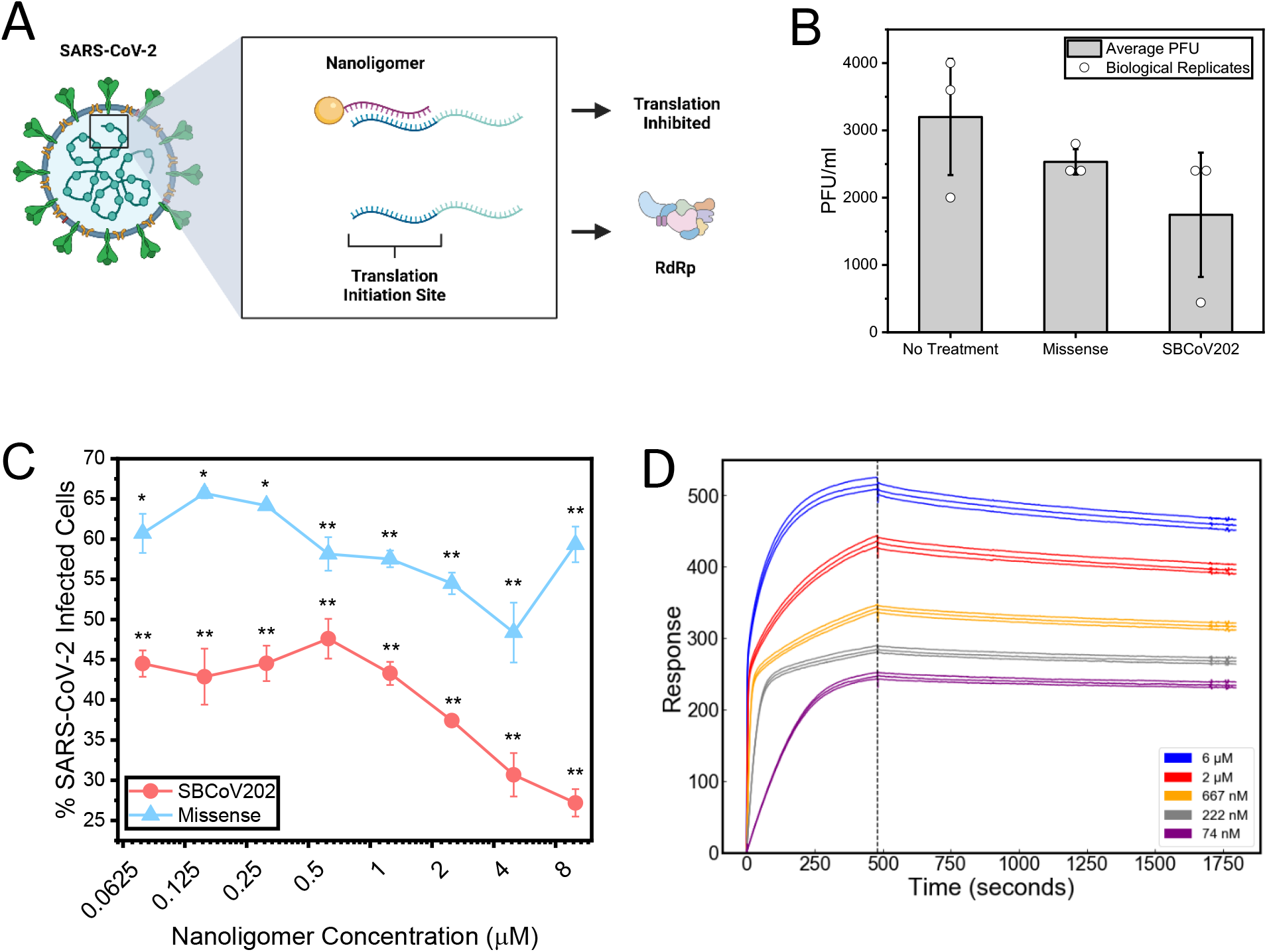
(**A**) Nanoligomers are PNA molecules conjugated to gold nanoparticles. Here, the Nanoligomer SBCoV202 is targeted to the translation initiation site of the RNA-dependent RNA polymerase within the SARS-CoV-2 genome. Blocking of this site through SBCoV202 binding is expected to prevent translation of RdRp, thus inhibiting viral replication. (**B**) A single dose of 10 μM of SBCoV202 is able to reduce plaque-forming units (PFU) of virus in infected Vero E6 cells, while missense Nanoligomer with PNA targeting no sequence in the viral or human circulating RNA do not result in a significant drop in viral load at this concentration. (**C**) Human A549-hACE2 lung epithelial cells infected with SARS-CoV-2, then fixed and immunostained for viral nucleocapsid protein, show lower percentages of infected cells when treated with SBCoV202 compared to missense at concentrations ranging from 0.078 to 10 μM. P-values were calculated with respect to three replicates of a no-treatment water control, shown in Figure S1 A. (**D**) SBCoV202 binds to the target viral genomic sequence with a K_D_ of 3.37 nM and an association rate constant (ka) of 18,390 M^-1^s^-1^, according to surface plasmon resonance measurements.

The Nanoligomer used here, SBCoV202, was tested for safety in a murine model. While animal species that are both susceptible to SARS-CoV-2 and are widely available in laboratory settings are limited,^34–36^ safety studies may be conducted similarly to any potential drug compound. As such, safety and biodistribution studies were performed using BALB/c mice. Measured parameters included overall body weight, as weight loss is a strong indicator of toxicity; inflammatory cytokine levels in the serum; histology of key organs; and assessment of biodistribution and clearance. The method of Nanoligomer administration to the mouse may also influence safety profiles. Therefore, three administration types were tested, namely intranasal (IN) administration, intravenous (IV) injection to the tail vein, and intraperitoneal (IP) injection. While IN administration was expected to provide the highest concentration to the infection site, the lung, and have the most clinical relevance,^37^ IV injection results in the highest concentration at key organs such as the liver and kidney, which allows for more thorough assessment of organ toxicity. IP injection loosely mimics an orally available drug and gives information on how first-pass organs may respond to the nanoligomers.^38^ We show here that the SBCoV202 Nanoligomer treatment is safe in mice and shows favorable biodistribution throughout the body, making it a strong candidate for a new antiviral treatment against COVID-19.

## Results

### Nanoligomer Performance In Vitro

The efficacy of SBCoV202 against SARS-CoV-2 and its binding affinity to the target sequence were assessed. Vero E6 cells infected with SARS-CoV-2, treated with 10 μM of SBCoV202 and assessed for viral load using a plaque assay 24 hours post infection and treatment, showed reduction in viral plaque forming units (PFU) (Figure 1B). A missense Nanoligomer targeting no predicted circulating miRNA, mRNA, or viral genome sequences was used as a negative control. This missense Nanoligomer also reduced the number of PFU per ml when compared with the no-treatment group, though to a lesser degree than SBCoV202.

The efficacy of SBCoV202 in inhibiting SARS-CoV-2 infection was assessed in human lung epithelial cells expressing human ACE2 (A549-hACE2) infected with SARS-CoV-2. Once treated with SBCoV202, cells were fixed and immunostained for the viral nucleocapsid (N) protein. SBCoV202 inhibited viral infection to a higher degree than missense Nanoligomers at concentrations ranging from 0.078 to 10 μM (Figure 1C). At 10 μM concentration, for example, Nanoligomers significantly reduced the percent of infected cells when compared to no treatment (p = 0.0002, Figures 1C and S1 A). Viral inhibition generally correlated with Nanoligomer concentration.

Viral abundance after SBCoV202 treatment was measured through viral mRNA transcriptional regulatory sequence N-protein (TRS-N) by quantitative reverse-transcription polymerase chain reaction (qRT-PCR) in infected A549-hACE2 cells treated with 10 μM SBCoV202 or missense Nanoligomers and normalized to abundance in cells treated with a water control (Figure S1 B). The TRS-N abundance is correlated with overall viral abundance, as the sequence is found in the genome of each viral particle.^39^ SBCoV202 treatment resulted in over a 7-fold reduction in viral mRNA abundance.

To quantify the binding affinity of SBCoV202 to the translation initiation site of RdRp, we performed surface plasmon resonance (SPR) measurements on different concentrations of Nanoligomers flowed over a DNA oligomer containing fragments of the viral genome with the translation initiation site sequence, complimentary to the SBCoV202 sequence. SBCoV202 binds to this site in anti-parallel orientation, the preferred orientation for strong PNA:DNA or PNA:RNA interactions.^40^ The binding response was measured during analyte injection (Figure 1D, up to vertical dashed line) and the resulting fit yielded an association rate constant (k_a_) of 18,390 M^-1^s^-1^. Subsequent dissociation was slow (dissociation rate constant k_d_ = 6.236×10^-5^ s^-1^), which is characteristic of PNA bound to nucleic acid targets.^41^ Using these results, we found that SBCoV202 strongly binds its complementary target with a measured dissociation constant K_D_ of 3.37 nM. We also performed SPR measurements on the missense Nanoligomer binding with the same DNA oligomer containing the RdRp translation initiation sequence. The response at different missense Nanoligomer concentrations was low (Figure S1 C), suggesting negligible levels of non-specific binding. These data provide *in vitro* evidence that Nanoligomers can strongly and specifically bind their complementary intended targets for long times.

### Nanoligomer Safety Assessment in Mice

Safety of SBCoV202 was first tested by administering mice with Nanoligomers, then keeping the mice under observation for 5 days before euthanizing and examining serum parameters and organ morphology. Three administration routes were assessed, with five mice tested for each experimental group. Intranasal (IN) administration and intravenous (IV) injection were tested at 0, 1, 2, 5, and 10 mg/kg body weight of SBCoV202. Intraperitoneal (IP) injection was tested at only 0 and 10 mg/kg body weight of SBCoV202.

Mouse body weight did not drop more than 1 g from Day 0 weight for all mice (Figures 2, S2, and S3), which falls within measurement error of our scale. Mice appeared healthy and active, with no noticeable changes to behavior or appearance as a result of the treatment. Albumin levels in the serum, which are an indicator of hepatic and renal health,^42,43^ were not significantly reduced in treatment groups compared to control groups administered with phosphate buffered saline (PBS) (Figures 3A and S4 A).

**Figure 2:**
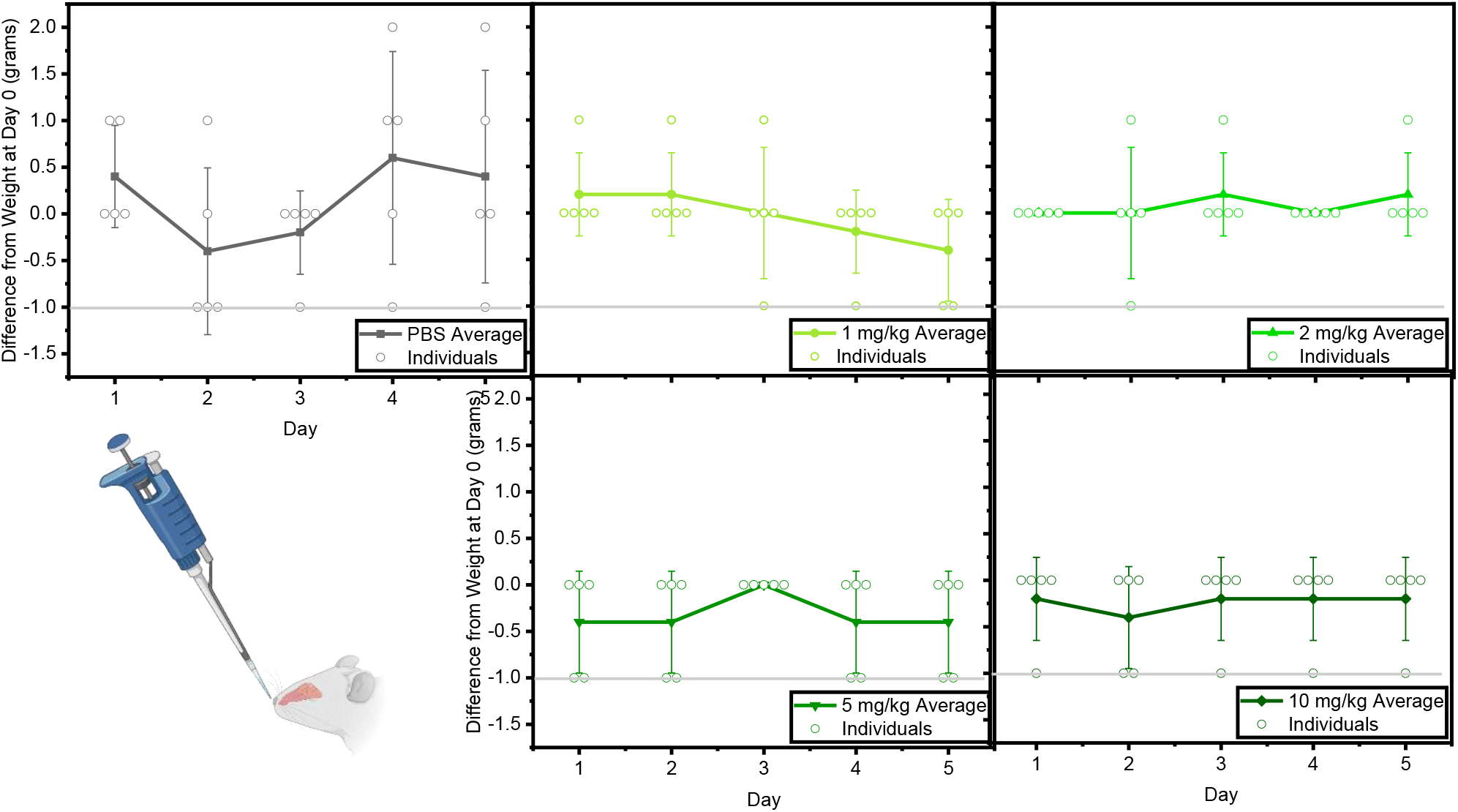
Intranasal administration of SBCoV202 did not cause weight loss in mice. Mice were administered with SBCoV202 intranasally (visualized in bottom left) and monitored for 5 days. Body weight did not drop more than 1 gram below starting weight for any mouse. The horizontal gray line is used to indicate 1 gram below the weight on Day 0 of the study. The p value was greater than 0.05 for all groups and timepoints compared to the PBS control group.

**Figure 3:**
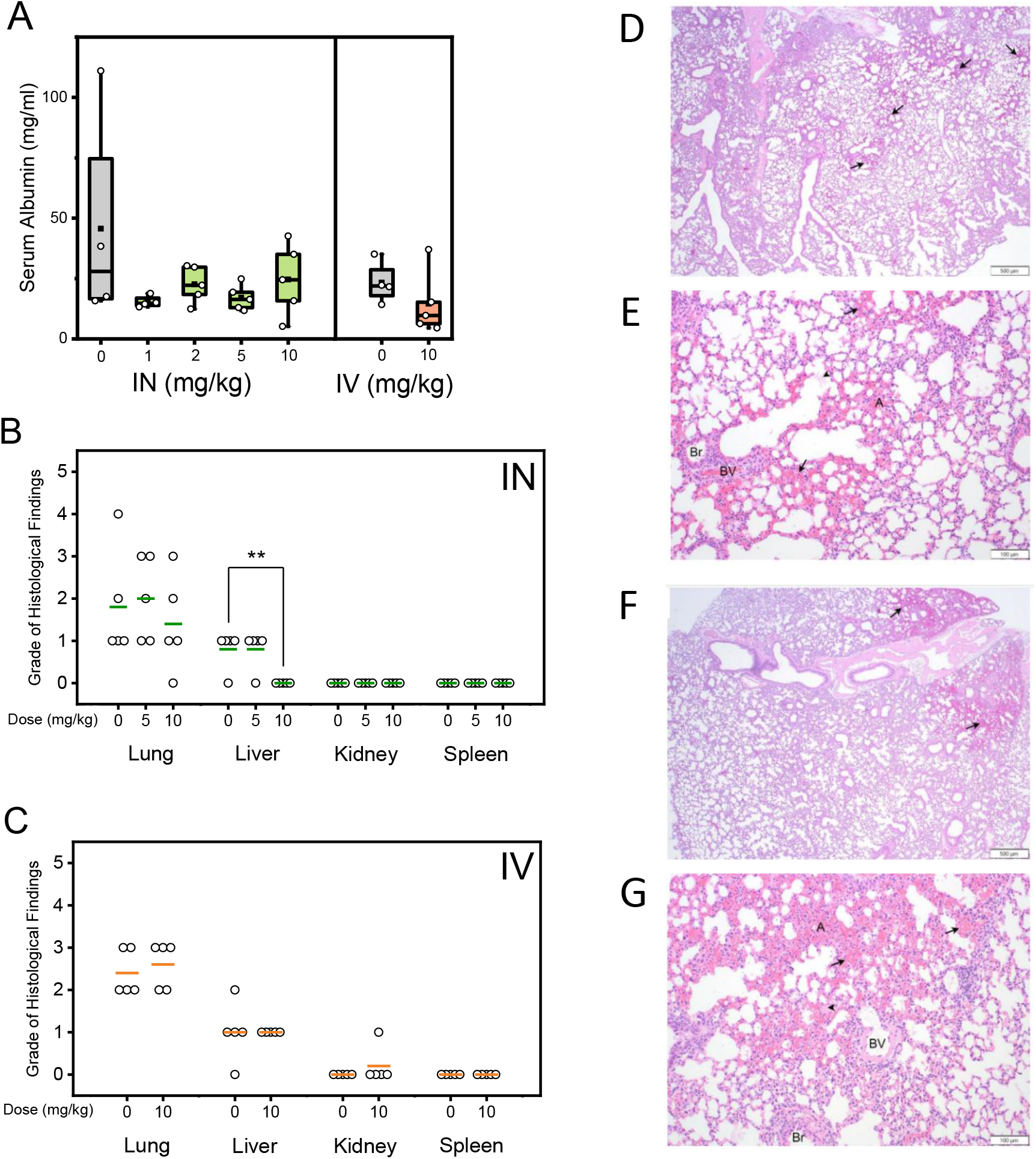
Administration of SBCoV202 did not result in significant changes to albumin levels in serum (**A**) or morphology of lung, liver, kidney, or spleen based on histology with H&E staining (**B-C**), regardless of administration method. Histology was graded on a 0-5 scale with 0=absent, 1=minimal, 2=mild, 3=moderate, 4=marked, 5=severe. Administration method is indicated in the upper right (**B-C**) corner of each plot and colored bars represent mean. ** indicates p < 0.01, otherwise p > 0.05. Similar multifocal regions of hemorrhage (arrows) are observed throughout the parenchyma in lungs of mice treated with PBS control (**D-E**) and SBCoV202 (**F-G**), often adjacent to blood vessels (BV), and characterized by free red blood cells within alveoli (A) and bronchioles (Br). Hemorrhage is sometimes accompanied by eosinophilic proteinaceous material (fibrin; arrowhead). Images taken at 20x (**D and F**) and 100x (**E and G**).

Histology and pathology were performed on fixed lung, liver, kidney, and spleen tissue. No difference was observed in organ morphology between PBS-treated control mice and mice receiving 5 or 10 mg/kg of SBCoV202, regardless of administration type (Figures 3 B-G and S4 B-F). While bleeding was observed in all IN-treated mice in the lungs, this hemorrhaging was confirmed to be due to the submandibular blood collection performed prior to euthanasia, rather than the Nanoligomer treatment, as the severity of hemorrhaging was comparable between treatment and control groups (Figure 3 D-G). Hemorrhaging observed in lungs of mice treated via IV or IP injection resulted from CO_2_ exposure during euthanasia, as PBS groups in these administration routes also showed similar bleeding (Figure S4 B-F).

Serum levels of two inflammatory markers, tumor necrosis factor α (TNF-α) and interleukin 6 (IL-6), were measured by enzyme-linked immunosorbent assay (ELISA). Levels for all treatment groups fell below the detection level of the assay (Table S1), which is expected for healthy and normal mice. In a separate study, mice were administered SBCoV202 before being euthanized at 1, 3, 6, or 24 hours after administration. In these mice as well, TNF-α and IL-6 levels remained below the level of detection (Table S2), indicating no acute inflammatory response.

Host immune response was further assessed by running serum from the IV and IP-injected mice in the 24-hour study through a 36-plex cytokine/chemokine panel (Figures 4, S5-S9). Only levels of IL-18 and IL-23 were higher in some treatment groups compared to the PBS control. As levels of other cytokines related to acute immune response, including IL-1β, IL-6, and TNF-α, were below the LOD, SBCoV202 is not shown to result in a measurable innate immune response. Further, levels of cytokines related to adaptive immune response and T-cell activation, including IFN-α, IFN-γ, IL-2, and IL-15, were also below LOD. While the increase in IL-18 indicates a possible T-cell response, the lack of downstream effects, as expected in IFN-γ levels, indicates that adaptive immune response is also minimal. SBCoV202, as a single injection at 10 mg/kg, therefore was not observed to be immunogenic.

**Figure 4:**
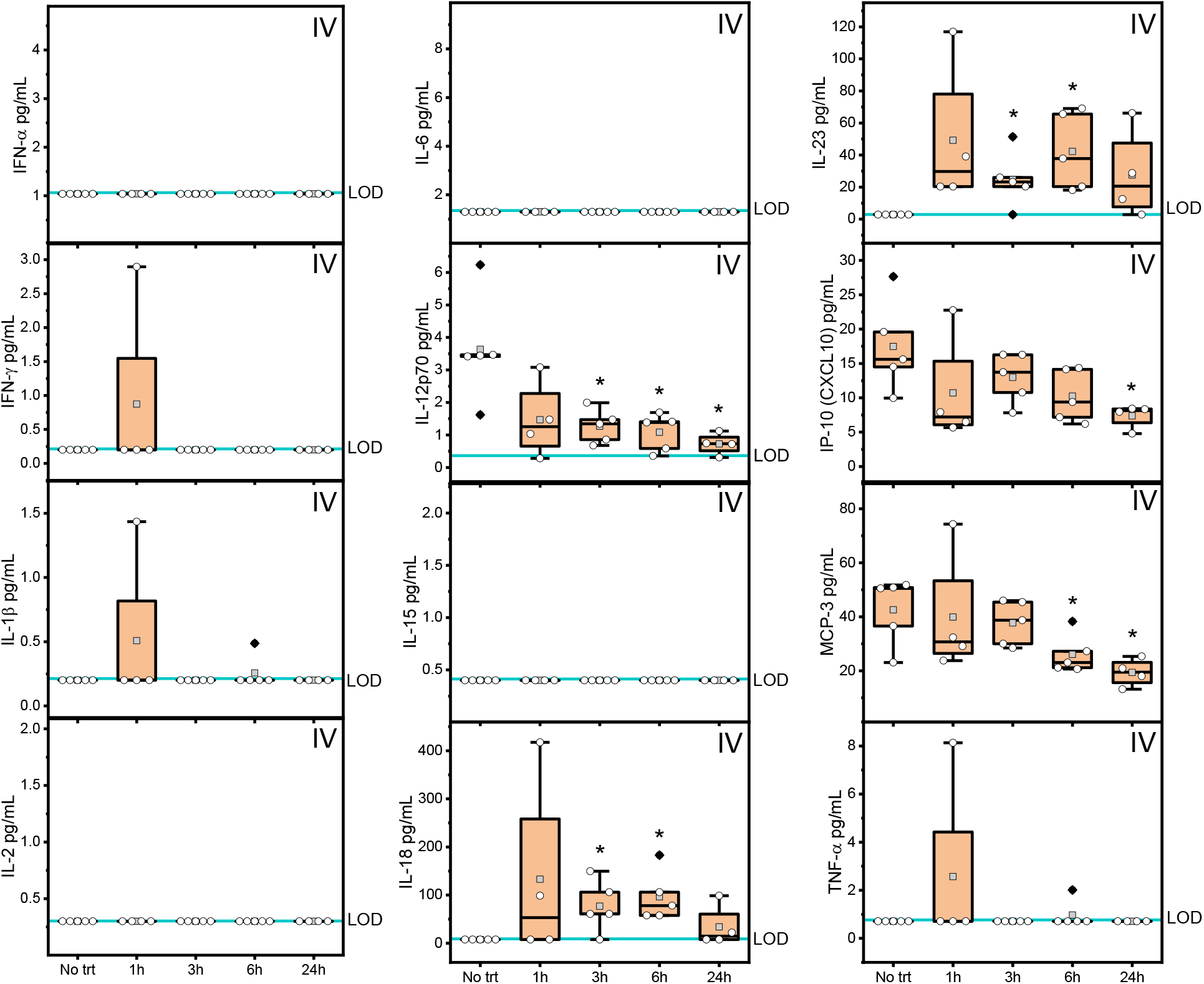
Key measurements from a 36-cytokine/chemokine panel performed on serum samples from mice treated with SBCoV202 via intravenous (IV) injection showed key cytokines largely below the limit of detection (LOD) of the assay, namely IFN-α, IFN-γ, IL-1β, IL-2, IL-6, IL-15, and TNF-α. The LOD is represented with a blue line. Levels of IL-12p70, IP-10, and MCP-3 were lower than the PBS control group at some timepoints. Only IL-18 and IL-23 levels were significantly increased compared to the control. * indicates p < 0.05. All other groups showed p > 0.05 compared to the PBS control group.

### Nanoligomer Biodistribution in Mice

Biodistribution within the first 24 hours after administration was measured in mice by measuring gold presence in organs due to Nanoligomer from mice euthanized 1, 3, 6, or 24 hours after a 10 mg/kg SBCoV202 administration. Gold was measured using inductively-coupled plasma mass spectrometry (ICP-MS), then converted to amount of SBCoV202 present in ng per g tissue using the mass ratio of gold per SBCoV202 molecule, or to nM using tissue densities.^44^

In IN-treated mice, the lungs showed high amounts of SBCoV202 in the first hour, and levels rapidly decreased in subsequent hours (Figure 5A). This indicates good bioavailability to the infection site. Even at 24 hours after administration, levels remained well above the SBCoV202 K_D_ of 3.37 nM (Figure 1D). In IV- and IP-treated mice, the highest amount measured in the lung was lower, at 20,000 ng/g tissue compared to 70,000 ng/g tissue (Figures 6A and S10A). This confirms that IN administration allows for more direct application of the treatment to the respiratory system, which is often the site of initial infection for SARS-CoV-2. IP treatment resulted in a slight delay in the peak concentration, at 3 hours instead of 1 hour after administration. Nanoligomer levels of this magnitude are also well above the K_D_ of the compound.

**Figure 5:**
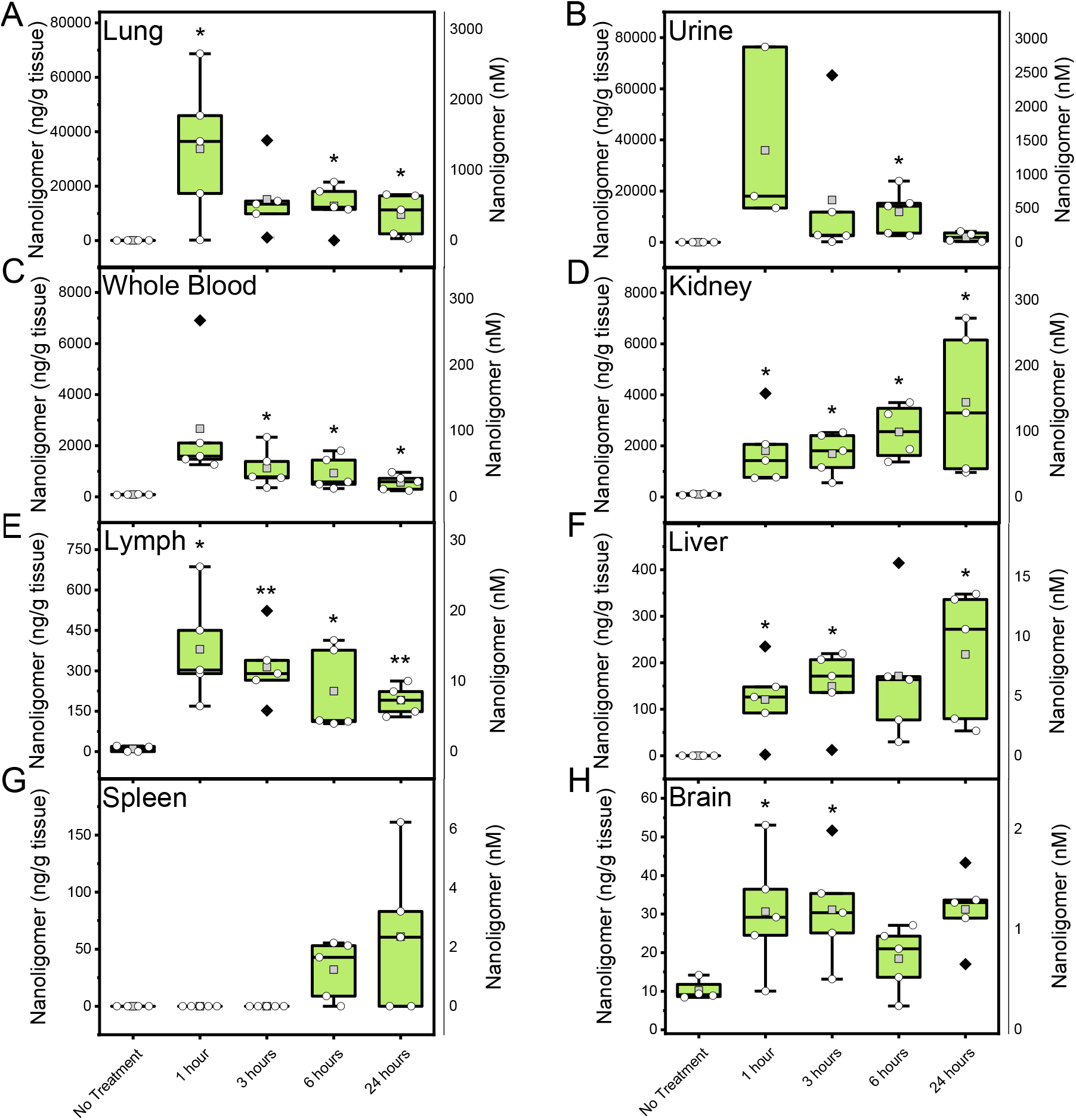
SBCoV202 biodistribution after intranasal administration showed SBCoV202 (shown as nanograms per gram of tissue on left, nanomolar on right) reaching the lungs (**A**), being cleared by the renal system (**D**), and excreted by the urinary system (**B**). SBCoV202 was circulated in the whole blood (**C**) and detected in lymph (**E**) and liver (**F**) tissue. Levels in the spleen (**G**) and brain (**H**) were lower than other organs. P-value indicated by * < 0.05, ** < 0.01, *** < 0.001, otherwise p > 0.05.

**Figure 6:**
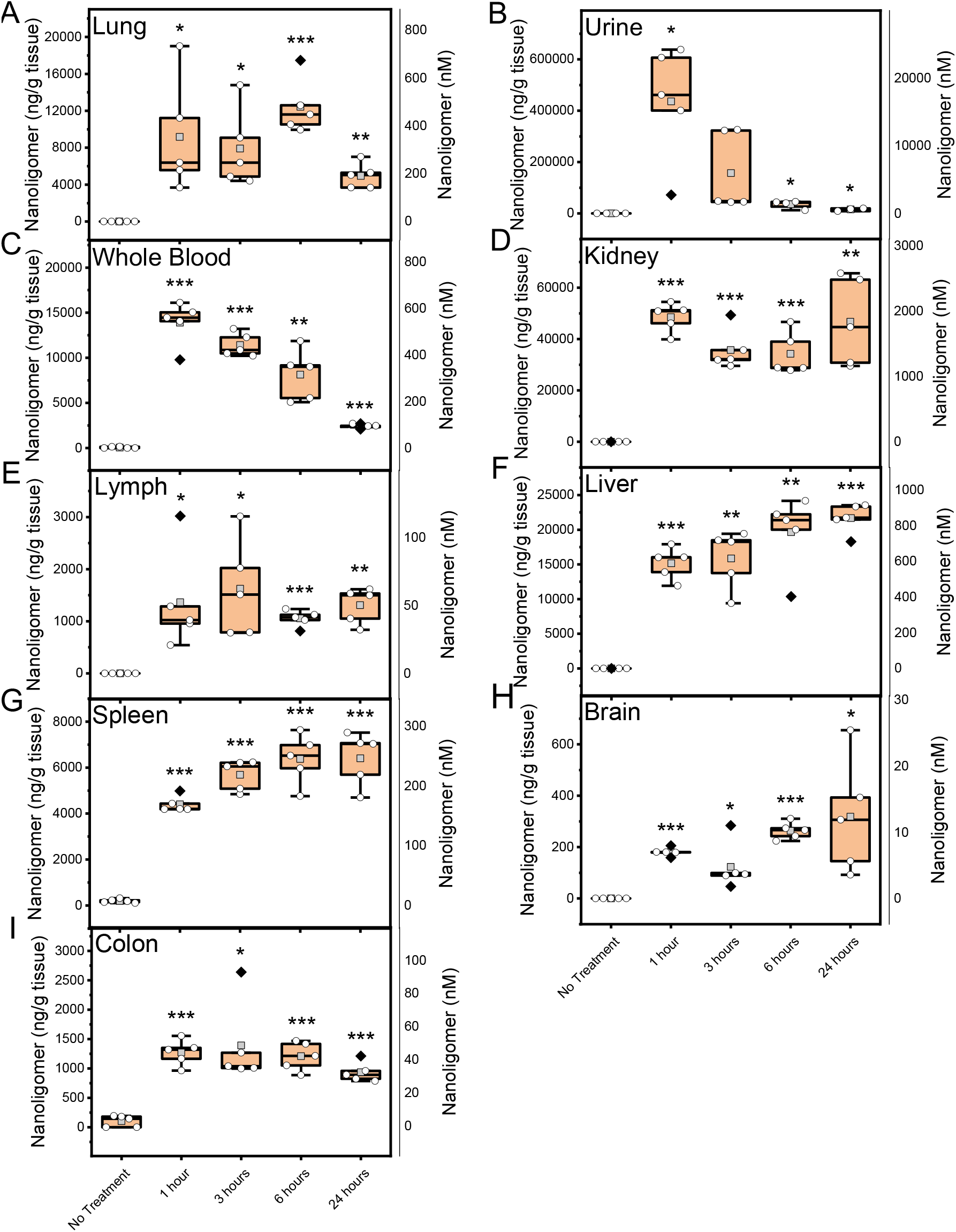
SBCoV202 biodistribution after intravenous administration showed SBCoV202 (shown as nanograms per gram of tissue on left, nanomolar on right) reaching the lungs (**A**), being cleared by the renal system (**D**), and excreted by the urinary system (**B**). SBCoV202 was circulated in the whole blood (**C**) and detected in lymph (**E**) and liver (**F**) tissue. Levels in the spleen (**G**), brain (**H**), and colon (**I**) were lower than other organs. P-value indicated by * < 0.05, ** < 0.01, *** < 0.001, otherwise p > 0.05.

SBCoV202 levels in urine indicate rapid urinary excretion. Peak levels in urine occur at 1 hour after administration in IN and IV-treated mice, but at 3 hours in IP-treated mice (Figures 5B, 6B, and S10 B). IP administration appears to result in slower biodistribution and clearance compared to the other two administration routes tested. The magnitude of SBCoV202 in urine was much lower in IN-treated mice, at 70,000 ng/g, compared to IV and IP, at 600,000 and 350,000 ng/g, respectively. In both IV and IP administrations, the peak level in urine was over an order of magnitude higher than peaks in any other organ or fluid. For treatment of infections in lung especially, IN administration maximizes the concentration of Nanoligomer available to fight infection. However, the high levels in urine observed in IV and IP groups are promising for rapid clearance with no accumulation.

Levels in the whole blood were approximately an order of magnitude lower than that observed in the lung and urine (Figure 5C) in IN groups. As expected, IV injection resulted in the highest peak concentration in the blood, at around 15,000 ng/g, but this value dropped rapidly with time (Figure 6C). In IP-treated mice, however, levels in blood stayed consistent for the first three timepoints, at around 13,000 ng/g, and only dropped at 24 hours (Figure S10 C). This offers IP administration as a favorable route for infections that have spread throughout the body, for sepsis, or for infections that require longer exposure or dosage interval of treatment.

In the kidney, levels of SBCoV202 rose over time in IN- and IP-treated mice (Figures 5D and 6D), but stayed more consistent in IV-treated mice (Figure S10 D). Levels in IN-treated mice were considerably lower than those in the other two administration routes, at nearly 7,000 ng/g tissue compared to up to 60,000 ng/g tissue. There is strong indication that the Nanoligomers are renally cleared prior to urinary excretion, with little involvement of other metabolic organs, considering the higher concentrations in kidney compared to other organs.

Levels in the liver were quite low in IN-treated mice compared to IV- and IP-treated mice, at < 500 ng/g tissue compared to 20,000 ng/g tissue or more (Figures 5F, 6F, and S10 F). Some SBCoV202 reached the lymph nodes and spleen (Figures 5E and G, 6E and G, and S10 E and G), though amounts were low relative to the kidney and liver. The lower organ levels for IN administration is likely due in part to the offset of higher concentrations in the lungs, but also due to inconsistent Nanoligomer amounts entering the body, as is commonly observed with IN administration.^37,45,46^ Amounts in the liver and spleen rose with time, indicating the pharmacokinetics of SBCoV202 biodistribution being on the scale of hours to reach these organs. Conversely, levels in the lymph dropped with time, which is in line with the function of the lymph nodes to act as a front-line defense against foreign materials in the body.

Over 100 ng/g tissue of SBCoV202 was observed in some IV and IP brain samples, and even in IN-treated mice (Figures 5H, 6H, and S10 H), SBCoV202 consistently reached the brain. This was unexpected considering typical inability of antisense therapeutics to cross the blood brain barrier unaided.^47,48^ The ability of the Nanoligomer to reach brain tissue, albeit at a much lower concentration than other organs, offers exciting new possibilities for applications in diseases affecting the brain.

Colons from IV- and IP-treated mice were also tested for SBCoV202 presence (Figures 6I and S10 I). Levels in the colon were consistently near 1,000 ng/g tissue for IV treatment. In IP-treated mice, initial levels were much higher at 6,000 ng/g tissue but dropped over time. This data does not give information on whether SBCoV202 was outside or within the colon, however, so further study would be necessary to gauge the potential of Nanoligomers in treating gastrointestinal infections via these administration routes.

At 5 days post-administration, organs from IN-treated mice were also assessed for gold via ICP-MS (Figure S11). This analysis shows that by 5 days post-treatment, levels of SBCoV202 in the organs is generally quite low, indicating that the compound was successfully cleared and no accumulation occurred. Some mice showed levels in organs above the K_D_ of the compound, where 90 ng/g tissue is approximately equivalent to the K_D_ of 3.37 nM, with slight variation depending on organ density. Only one or two mice per group showed SBCoV202 levels that could indicate widespread binding, in the urine, liver, spleen, (Figure S11 B) lungs, and kidneys (Figure S11 C).

## Discussion

SARS-CoV-2 spread rapidly around the world in the first half of 2020 and leading into present times. Further, the rapid emergence of variants of concern such as Delta and Omicron, the latter of which caused difficulties with vaccine immunity without booster shots,^5^ has left drug developers scrambling to maintain pace manufacturing effective strategies to counter disease spread. The Nanoligomer treatment presented here represents a nontoxic alternative to currently available treatments for COVID-19. The sequence-specific PNA is able to bind only the target region of the viral genome, without risk of off-target binding to human genetic material. SBCoV202 was demonstrated to be nontoxic to mice, causing no loss of weight or changes to serum parameters that would indicate adverse side effects. Lung, liver, kidney, and spleen tissue showed no damage as a result of SBCoV202 administration. Various cytokines and chemokines that would indicate innate and adaptive immune response were not elevated. Further, SBCoV202 was distributed throughout the body relatively evenly, with clear signs of urinary excretion occurring largely within a couple hours of administration.

The combination of organ histology showing no signs of damage and the healthy albumin levels in the serum indicate renal and hepatic health. While neutral antisense molecules like PNA typically demonstrate even distribution throughout the body,^47,49^ excepting the central nervous system,^47,48^ and rapid clearance from the blood via urinary excretion, the liver and kidney would be expected to show higher concentrations than other organs.^47,50,51^ This was generally true even with the addition of the gold nanoparticle conjugate, but while other studies show large fractions of over 70% of administered ASOs localizing to the liver,^47^ the Nanoligomers spread more evenly among the body’s organs, with biodistribution to the kidneys, lungs, lymph, and eventually even spleen. The higher SBCoV202 concentration in the kidney and liver at 24 hours compared to other organs did not seem to cause toxicity within the mice, which is in line with the expected nontoxic nature of naked PNA^29^ and the biocompatibility of gold in the body.^52^ SBCoV202 was able to reach other organs like lungs, colon, and even brain. While a higher fraction of total SBCoV202 was observed in the kidney than other organs, biodistribution throughout the body indicates that infection treatment could be efficacious at a variety of infection sites. In all administration types, for example, SBCoV202 concentrations were well above the compound K_D_ in the lungs, lymph, and whole blood. Further, with IP and IV-administered mice, concentrations of SBCoV202 were generally above the K_D_ of the compound in the liver, spleen, brain, and colon within 24 hours after administration.

Direct-acting ASO therapeutics offer a unique approach to infection treatment due to their high specificity. The development of direct-acting antivirals to improve patient outcomes has occurred previously in hepatitis C treatment. With hepatitis C, the introduction of direct-acting antivirals (DAA) led to an increase in sustained virologic response from 55% to as high as 99%.^26^ However, these compounds, which act on the virus by inhibiting key proteins, stalling RNA synthesis, or blocking nucleosides from binding polymerases,^10,53^ soon drove the emergence of drug resistance in hepatitis C virus.^26^ With ASO therapies like Nanoligomers, however, resistance can be circumvented due to the adaptability of the nucleic acid sequence. Even as the virus evolves and mutates, the Nanoligomer can also be adapted to counter resistant variants. This has become increasingly relevant in the COVID-19 pandemic, where emergence of new variants of concern such as Delta and Omicron have strained the healthcare system and threatened the efficacy of vaccines.^5^

Current antiviral treatments include nucleoside inhibitors and non-nucleoside inhibitors. Over two dozen nucleoside inhibitors have been developed, but few have gone beyond Phase III clinical trials.^13,53^ Of these, remdesivir became the first FDA-approved COVID-19 treatment. While generally deemed safe based on clinical trials thus far,^8,11,12^ remdesivir can cause adverse events including respiratory failure, low albumin or potassium, or low blood cell counts for red blood cells and platelets, in addition to milder side effects such as nausea.^12,14^ Remdesivir is also currently only approved for hospitalized patients with COVID-19, and is not recommended for use in those with impaired liver or kidney function due to risk of complications such as liver inflammation or accumulation and toxicity.^54^ Most non-nucleoside inhibitors targeting the RdRp generally bind allosteric sites on the RdRp to inhibit replication.^53^ Non-nucleoside inhibitor development has largely focused on the treatment of hepatitis C.^53^ SBCoV202 is unique from both these broad categorizations of antivirals, as it binds the viral genome to prevent the RdRp from being translated initially. While allosteric inhibitors rely on conservation of the specific binding site environment, Nanoligomers are more adaptable. Were a resistant variant to emerge, a simple genomic sequencing of the virus would allow for identification of the mutation preventing PNA binding, and a new complementary sequence could be synthesized and applied instead.

Nanoligomers offer advantages over other antisense oligonucleotide (ASO) platforms due to their lack of immunogenicity, even biodistribution, and potential for reaching the lymph and brain. SBCoV202 did not result in any upregulation of cytokines related to innate or acquired immune response. While a longer-term study would be necessary to confirm a lack of T cell response and antibody production, there is thus far no indication that Nanoligomers cause activity in the immune system (Figures 4, S5-S9). This is in contrast with siRNA therapies, PNA conjugated to CPP, and lipid-encapsulated ASOs.^29,31–33^ Nanoligomers are readily available to key organs such as the lungs, kidneys, liver, and spleen. They distribute evenly throughout the body without accumulating in any one organ. Clearance and excretion are rapid, leading to favorable pharmacokinetics. Further, the Nanoligomers show promise in being able to reach the lymph nodes and brain at efficacious concentrations, furthering potential applications of the treatment. Other ASOs have especially demonstrated difficulty in crossing the blood-brain barrier.^47,48^

SBCoV202 will require further testing before entering clinical trials. Pre-clinical testing in line with Good Laboratory Practices (GLP) that include larger sample size; confirmation in another species; testing of the effects on other bodily systems such as the cardiovascular system, central nervous system, and reproductive systems; urinalysis; and full clinical chemistry will be necessary to comprehensively characterize the effects of Nanoligomers in the body. The maximum tolerable dose is also unknown as of yet; the highest dose tested here of 10 mg/kg showed no deleterious effects, indicating higher doses may also be safe. Efficacy studies in infected animals will be performed to determine the therapeutic window of the Nanoligomers and to ensure that the mechanism of action, namely prevention of translation of the RdRp, is adequate to inhibit viral replication and decrease viral load in infected animals.

## Conclusions

The Nanoligomer treatment SBCoV202 was nontoxic in mice and showed favorable biodistribution, making it an attractive candidate as an antiviral for treatment of COVID-19. The compound has a PNA sequence that can be adapted to circumvent resistance emergence and was able to inhibit viral spread *in vitro*. No adverse response was observed in mice treated with the SBCoV202, whether through intranasal, intravenous, or intraperitoneal administration. Treatment with SBCoV202 did not result in unintended immune response or organ damage and had no effect on body weight or mouse behavior. Distribution of the compound was relatively even throughout body organs, with intranasal administration allowing for higher concentrations to the lungs, where the infection is most severe in the majority of patients. Biodistribution to other key organs also resulted in Nanoligomer concentrations above the K_D_ of SBCoV202, indicating promise for efficacy throughout the body. With further studies on the pharmacokinetics, maximum safe dose, and *in vivo* efficacy in infected animals, SBCoV202 may offer a novel alternative antiviral treatment during the global COVID-19 pandemic.

## Materials and Methods

### Nanoligomer Design and Synthesis

Nanoligomers were designed and synthesized at Sachi Bioworks (Boulder, CO, USA) according to the methods described in McDonald et al.^55^ After identification of the target translation initiation site, a PNA sequence complementary to the target was designed. The FAST platform^55–57^ allows for the scanning of the human genome for the appearance of similar sequences, which could result in off-target binding and side effects from compound administration. Of all potential target sequences within the translation initiation site, SBCoV202 was built from the sequence exhibiting the highest solubility at biologically relevant pH and lowest incidence of self-complementing sequences and off-targets within the human genome.^55^ Following synthesis of the 15-base pair PNA molecules via solid-phase Fmoc chemistry on an Apex 396 peptide synthesizer (AAPPTec, LLC), the PNA were conjugated to gold nanoparticles to form Nanoligomers. Nanoligomers were purified by size-exclusion filtration. Conjugation and concentration were confirmed using absorbance measurements for detection of PNA and quantification of gold nanoparticles.

### Surface Plasmon Resonance Measurements

A DNA oligomer containing 30 nucleotides of the human genome (hg38) with the RdRp translation initiation site binding target was biotinylated on the 5’ end (IDT). This oligomer probe was diluted to 5 μg/mL using 1x HBS-EP+ buffer. This 1x HBS-EP+ buffer was also used as running buffer during the SPR measurements. These oligomers were attached to a streptavidin-coated flow cell (Cytiva). The chip was first cleaned using a 10 μL injection of 20 mM NaOH/1M NaCl solution. The probe was subsequently injected onto a flow cell of the chip. RU responses of 256 were observed, indicating probe binding to the chip. One flow cell on the chip was used as a reference; while it was cleaned as described above, no probe was attached.

SPR measurements were carried out in accordance with previously established protocols.^41,58^ The measurements were carried out on a Biacore 3000 instrument (Cytiva). Nanoligomer analyte was serially diluted using 2-fold dilutions from 74 nM-6 μM. To measure association, 90 μL of Nanoligomer was run over the chip for 3 minutes at a flow rate of 30 μL/min. Dissociation was subsequently measured for 30 minutes. The chip surface was regenerated with a 10 μL injection of 20 mM NaOH+1M NaCl solution. Measurements for each concentration were repeated in triplicate. Data from the SPR measurements was analyzed using Scrubber 2.0 (BioLogic).

### Virus Propagation and Plaque Assay

SARS-related coronavirus 2 (SARS-CoV-2), Isolate USA-WA1/2020 (NR-52281) was deposited by the Center for Disease Control and Prevention and obtained through BEI Resources, NIAID, NIH. SARS-CoV-2 was propagated in Vero E6 cells in DMEM supplemented with 2% FBS, 4.5 g/L D-glucose, 4 mM L-glutamine, 10 mM Non-Essential Amino Acids, 1 mM Sodium Pyruvate and 10 mM HEPES. Infectious titers of SARS-CoV-2 were determined by plaque assay in Vero E6 cells in Minimum Essential Media supplemented with 2% FBS, 4 mM L-glutamine, 0.2% BSA, 10 mM HEPES and 0.12% NaHCO3 and 0.7% agar. 48 hours after addition of plaque overlay, cells were fixed with 5% formaldehyde for 24 hours. After fixation overlay was removed, cells were stained with 0.2% Crystal Violet (in 20% EtOH) and plaques were counted.

### Quantitative Reverse-Transcription PCR (qRT-PCR) of Viral RNA

qRT-PCR of SARS-Cov-2 RNA was performed as previously described.^59^ RNA was extracted from human lung epithelial cells expressing human ACE2 (A549-hACE2) grown in 96-well plates by using the RNeasy 96 Kit (Qiagen) per the manufacturer’s instructions. RNA was reverse-transcribed and PCR amplified using Luna Universal One-Step RT-PCR Kit (NEB). SARS-CoV-2 replication was assessed by using primers specific to the N mRNA (Forward 5’-CTCTTGTAGATCTGTTCTCTAAACGAAC-3’; Reverse 5’-GGTCCACCAAACGTAATGCG-3’). SARS-CoV-2 N mRNA levels were normalized to beta tubulin (Forward 5’-GCCTGGACCACAAGTTTGAC-3; Reverse 5’-TGAAATTCTGGGAGCATGAC-3’). Reactions were ran and analyzed on a Lightcycler 480 II Instrument (Roche). Relative quantification was calculated by comparing the cycle threshold (*C_t_*) values using ΔΔC_*t*_. Significance was determined using a two-tailed unpaired Student’s *t*-test.

### Immunofluorescence of Nucleocapsid (N) Protein

Quantification of viral infection was performed as previously described.^59^ Briefly, at indicated times after infection A549-hACE2 cells were fixed with 5% formaldehyde and immunostained for nucleocapsid (N) protein. N protein was visualized with a secondary antibody labeled with AlexaFlour-488 (ThermoFisher). SARS-CoV-2 nucleocapsid (N) antibody (clone 1C7C7) was obtained from the Center for Therapeutic Antibody Discovery at the Icahn School of Medicine at Mount Sinai. Nuclei were stained with DAPI. Full wells were imaged and quantified for SARS-CoV-2 infected cells using a Celigo imaging cytometer (Nexcelom Biosciences). All infections with SARS-CoV-2 were performed with 3 biological replicates.

### Five-Day Safety Studies

Mouse studies were conducted in accordance with the University of Colorado IACUC approval under protocol #2807. Female mice of age 8-12 weeks, purchased from Envigo (Indianapolis, IN) were divided into groups of five to receive either 1, 2, 5, or 10 mg/kg of SBCoV202, diluted in sterile PBS. A control group was administered with sterile PBS of the same volume. Intranasal administration was achieved by anesthetizing the mouse with isoflurane, holding the mouse at a 45° angle, and gently pipetting 25 μl of SBCoV202 solution into the nostrils.^45^ Intravenous initial drug administration was performed by injecting 100 μl in the lateral tail vein of the mouse. Intraperitoneal drug administration was performed by injecting 100 μl in the intraperitoneal space of the mouse on the right side of the lower abdomen. Only 10 mg/kg of SBCoV202 was tested for safety in IP administration. Mice were then monitored for five days before being euthanized. IN group mice were euthanized by cervical dislocation. Cervical dislocation was chosen to preserve the integrity of lung tissue, which may be damaged by carbon dioxide inhalation euthanasia. Prior to euthanasia, blood samples were collected via submandibular blood draw, wherein a lancet is used to pierce the facial vein and blood is allowed to drip into a collection vessel, and urine samples were collected by palpating the bladder. Euthanasia was performed using CO2 exposure for IV- and IP-treated mice, followed by blood collection via intracardial draw. Urine samples were collected by euthanizing each mouse separately in a clean and empty cage and collecting urine released from the bladder upon death. Each day of the study, mouse weight was measured. Once the mouse was euthanized, samples were collected of brain, lymph node, lung, liver, spleen, and kidney tissue. Colon was also collected from IV- and IP-treated mice. Collected tissues were rinsed in PBS and either drop-fixed in 10% formalin for histology or immediately frozen at −80 °C for ICP-MS.

Blood samples incubated on ice for one hour to clot before being centrifuged at 10,000 g for 10 min to collect serum. Serum was frozen at −80 °C until use in ELISAs. ELISAs for TNF-α and IL-6 (Duoset ELISA, R&D Systems, Minneapolis, MN) and albumin (Immunology Consultants Laboratory, Inc., Portland, OR) were used according to manufacturer directions, with serum diluted 1:100 for TNF-α and IL-6 and 1:500,000 for albumin, and plates were measured at 450 nm in a SpectraMax iD3 plate reader (Molecular Devices, LLC, San Jose, CA) using SoftMax Pro software. Standard curves were built using 4-parameter logistic regression.^60^

Fixed samples were removed to 70% ethanol after 24 hours. Histology was performed by Inotiv (Boulder, CO). Formalin-fixed liver, spleen, kidney, and lung samples from mice that received either 10 mg/kg of SBCoV202 or PBS control were processed. Tissues were blocked in paraffin. One slide per block was sectioned at 4 μm and stained with hematoxylin and eosin (H&E). Glass slides were evaluated by an ACVP-board-certified veterinary pathologist, using light microscopy. Histologic findings in each tissue were diagnosed and graded for severity on a 0-5 scale; specifically in this study, alveolar hemorrhage was graded 0=absent, 1=minimal (<10% of lung affected), 2=mild (10-25% of lung affected), 3=moderate (26-50% of lung affected), 4=marked (51-75% of lung affected), 5=severe (>75% of lung affected). This scale was also used to assess inflammation in the kidney (mononuclear and neutrophilic, focal, capsule, and with fibrosis) and liver (subacute, multifocal, and random). No histological findings were observed in any spleen samples.

Samples for ICP-MS were later thawed and homogenized in a TissueLyser II bead mill (Qiagen, Germantown, MD) at 30 Hz for 3 min with the addition of 1 μl deionized water per 1 mg organ tissue. Volumes of homogenates ranging from 10 to 130 μl were then digested in 500 μl of aqua regia (3:1 hydrochloric acid to nitric acid) for 4 hours at 100 °C. Pellets were resuspended in water and analyzed with a NexION 2000B single quadropole ICP-MS (PerkinElmer, Waltham, MA). A Meinhard nebulizer was used with a cyclonic glass spray chamber for the introduction of the sample. A nickel sample and skimmer cone were used with an aluminum hyperskimmer cone. The ICP-MS was optimized daily with a calibration solution of 1 ppb In, Ce, Be, U, and Pb. Data was collected using the sample acquisition module in Syngistix software (version 2.3).^197^ Au was the analyte monitored. A seven-point linear standard curve was generated at a concentration range of 0-250 ppb. Indium was spiked into each sample at a concentration of 5 ppb. Linearity of the standard curve was defined at an R^2^ value of greater than 0.995. The 1000 ppm gold standard solution was purchased from Ricca Chemical Company (Arlington, TX). TraceMetal grade 70% HNO3 was purchased from ThermoFisher Scientific (Waltham, MA). All H2O used was from a Milli-Q system (Millipore, Burlington, MA). Data was analyzed using Microsoft Excel by converting measured gold concentrations to ppb (ng/g tissue) via organ homogenate volume used. This ppb value was then converted to moles of gold, then moles of SBCoV202, per gram of tissue, and finally to ng of SBCoV202 per g of tissue using the molecular weight of the SBCoV202 molecule (27,775.35 g/mol). This was done to provide a clear representation of data consistent with antisense therapeutic data reported in other studies.^47,49,50,61,62^ The final conversion factor was 6.4087.

### Twenty-Four Hour Biodistribution Studies

Female mice of age 8-12 weeks (Envigo, Indianapolis, IN) were divided into groups of five. Each mouse received 10 mg/kg SBCoV202 either via IN, IV, or IP administration. Blood and urine samples were collected prior to cervical dislocation at 1, 3, 6, or 24 hours after SBCoV202 administration for IN mice, or immediately after CO_2_ exposure for IV and IP mice. A small aliquot of 30 μl of whole blood was set aside in a heparin-coated tube for ICP-MS analysis; the remaining blood was kept in a standard tube to clot for serum retrieval. A control group of four mice that received no treatment was also euthanized by cervical dislocation for comparison to IN mice. While a matching sample size of 5 was intended, one mouse arrived from the vendor deceased, and as such the control group was limited to 4 animals. A second control group of 5 mice was euthanized by CO_2_ exposure for comparison to IV and IP groups. Organ samples were collected and analyzed as described above, however no samples were drop-fixed in formalin.

### Cytokine/Chemokine Panel for Measuring Immune Response

Serum from IV- and IP-treated mice in the biodistribution studies were additionally assessed in a 36-plex cytokine/chemokine panel. Quantification of cytokines was performed using 25 μL of serum in Invitrogen Cytokine & Chemokine Convenience 36-Plex Mouse ProcartaPlex Panel 1A (EPXR360-26092-901) following manufacturers instruction and was analyzed on a Luminex MAGPIX xMAP instrument. The panel measured levels of the following cytokines and chemokines: ENA-78 (CXCL5), Eotaxin (CCL11), GRO-α (CXCL1), IP-10 (CXCL10), MCP-1 (CCL2), MIP-1α (CCL3), MIP-1β (CCL4), MIP-2α (CXCL2), RANTES (CCL5), G-CSF (CSF-3), GM-CSF, IFN-α, IFN-γ, IL-1α, IL-1β, IL-2, IL-3, IL-4, IL-5, IL-6, IL-9, IL-10, IL-12p70, IL-13, IL-15/IL-15R, IL-17A (CTLA-8), IL-18, IL-22, IL-23, IL-27, IL-28, IL-31, LIF, MCP-3 (CCL7), M-CSF, and TNF-α. Known concentrations of each cytokine/chemokine were used to build standard curves, from which amounts could be calculated per sample.

### Pharmacokinetic Biodistribution Analysis

The PK was examined through analyzing data from the 24-hour biodistribution study, by converting moles of SBCoV202 amounts in different organ tissues to concentrations (mole/volume) by using tissue densities.^63^ Briefly, density of blood (1050±17 kg/m^3^), brain (1046±6 kg/m^3^), colon (1045 kg/m^3^), kidney (1066±56 kg/m^3^), liver (1079±53 kg/m^3^), lung (1050 kg/m^3^), lymph (1019 kg/m^3^), and spleen (1089±64 kg/m^3^) were used to calculate SBCoV202 concentrations in different organ tissues.

### Statistical Analysis of Data and Data Visualization

Unless otherwise stated, data was analyzed using Microsoft Excel. P-values for samples were calculated using a student’s two-tailed T-test, with significance defined as p < 0.05. Standard curves for ELISAs were generated using a four-parameter logistic regression MATLAB (MathWorks) code by Cardillo.^60^ Data from the SPR measurements was analyzed using Scrubber 2.0 (BioLogic). Figures 1A, 2 (bottom left), S2 (bottom left), and S3 (right) were created with BioRender (BioRender.com). Tables S1 and S2 were created with Microsoft Office. Figures 1D and S1 C were created with Jupyter (Project Jupyter). Figure 3 D-G and Figure S4 C-E were taken by Inotiv. All other figures were created with Origin (OriginLab).

## Author Contributions

The manuscript was written through contributions of all authors. C.R.M., C.M.C., and A.C. designed the experiments. C.R.M. and C.M.C. conducted 5-day safety studies and biodistribution studies. C.R.M. performed ELISAs and homogenized and digested samples for ICP-MS. C.M.C. and P.N. conducted multi-plex ELISAs on blood serum. N.J.O., and T.R.A synthesized PNA. P.N. synthesized Nanoligomers. N.J.O. performed SPR analysis. T.X.J. performed A549 cell *in vitro* studies and qPCR. K.L.R. and S.B. performed ICP-MS. C.R.M., C.M.C., N.J.O., P.N., and A.C. analyzed the experimental data. C.R.M., C.M.C., N.J.O., T.X.J., K.L.R., P.N., and A.C. wrote the manuscript. All authors discussed the results and edited the manuscript. All authors have given approval to the final version of the manuscript.

## Acknowledgments

We acknowledge L. Larson and J. Figueroa Hernandez for assistance with mouse procedures and guidance on IACUC protocols. We acknowledge J. Lindenberger for developing SPR protocols and conducting SPR measurements.

## Supporting Information

Supporting information is provided as a separate PDF file and includes Supplemental Tables S1-S2 and Supplemental Figures S1-S11.

## References

(1) Borio, L. L.; Bright, R. A.; Emanuel, E. J. A National Strategy for COVID-19 Medical Countermeasures. J. Am. Med. Assoc. 2022, Online, E1–E2. https://doi.org/10.1056/nejme2034495.

(2) Sadoff, J.; Gray, G.; Vandebosch, A.; Cárdenas, V.; Shukarev, G.; Grinsztejn, B.; Goepfert, P. A.; Truyers, C.; Fennema, H.; Spiessens, B.; Offergeld, K.; Scheper, G.; Taylor, K. L.; Robb, M. L.; Treanor, J.; Barouch, D. H.; Stoddard, J.; Ryser, M. F.; Marovich, M. A.; Neuzil, K. M.; Corey, L.; Cauwenberghs, N.; Tanner, T.; Hardt, K.; Ruiz-Guiñazú, J.; Le Gars, M.; Schuitemaker, H.; Van Hoof, J.; Struyf, F.; Douoguih, M. Safety and Efficacy of Single-Dose Ad26.COV2.S Vaccine against Covid-19. N. Engl. J. Med. 2021, 384 (23), 2187–2201. https://doi.org/10.1056/nejmoa2101544.

(3) Polack, F. P.; Thomas, S. J.; Kitchin, N.; Absalon, J.; Gurtman, A.; Lockhart, S.; Perez, J. L.; Pérez Marc, G.; Moreira, E. D.; Zerbini, C.; Bailey, R.; Swanson, K. A.; Roychoudhury, S.; Koury, K.; Li, P.; Kalina, W. V.; Cooper, D.; Frenck, R. W.; Hammitt, L. L.; Türeci, Ö.; Nell, H.; Schaefer, A.; Ünal, S.; Tresnan, D. B.; Mather, S.; Dormitzer, P. R.; Şahin, U.; Jansen, K. U.; Gruber, W. C. Safety and Efficacy of the BNT162b2 MRNA Covid-19 Vaccine. N. Engl. J. Med. 2020, 383 (27), 2603–2615. https://doi.org/10.1056/nejmoa2034577.

(4) Baden, L. R.; El Sahly, H. M.; Essink, B.; Kotloff, K.; Frey, S.; Novak, R.; Diemert, D.; Spector, S. A.; Rouphael, N.; Creech, C. B.; McGettigan, J.; Khetan, S.; Segall, N.; Solis, J.; Brosz, A.; Fierro, C.; Schwartz, H.; Neuzil, K.; Corey, L.; Gilbert, P.; Janes, H.; Follmann, D.; Marovich, M.; Mascola, J.; Polakowski, L.; Ledgerwood, J.; Graham, B. S.; Bennett, H.; Pajon, R.; Knightly, C.; Leav, B.; Deng, W.; Zhou, H.; Han, S.; Ivarsson, M.; Miller, J.; Zaks, T. Efficacy and Safety of the MRNA-1273 SARS-CoV-2 Vaccine. N. Engl. J. Med. 2021, 384 (5), 403–416. https://doi.org/10.1056/nejmoa2035389.

(5) Garcia-Beltran, W. F.; St Denis, K. J.; Hoelzemer, A.; Lam, E. C.; Nitido, A. D.; Sheehan, M. L.; Berrios, C.; Ofoman, O.; Chang, C. C.; Hauser, B. M.; Feldman, J.; Gregory, D. J.; Poznansky, M. C.; Schmidt, A. G.; Iafrate, A. J.; Naranbhai, V.; Balazs, A. B. MRNA-Based COVID-19 Vaccine Boosters Induce Neutralizing Immunity against SARS-CoV-2 Omicron Variant. medRxiv Prepr. Serv. Heal. Sci. 2021, 1–10. https://doi.org/10.1101/2021.12.14.21267755.

(6) The Medical Letter. An EUA for Bamlanivimab - A Monoclonal Antibody for COVID-19. J. Am. Med. Assoc. 2021, 325 (9), 880–881. https://doi.org/10.1056/nejmoa2029849.

(7) Taylor, P. C.; Adams, A. C.; Hufford, M. M.; de la Torre, I.; Winthrop, K.; Gottlieb, R. L. Neutralizing Monoclonal Antibodies for Treatment of COVID-19. Nat. Rev. Immunol. 2021, 21 (6), 382–393. https://doi.org/10.1038/s41577-021-00542-x.

(8) Gavriatopoulou, M.; Ntanasis-Stathopoulos, I.; Korompoki, E.; Fotiou, D.; Migkou, M.; Tzanninis, I. G.; Psaltopoulou, T.; Kastritis, E.; Terpos, E.; Dimopoulos, M. A. Emerging Treatment Strategies for COVID-19 Infection. Clin. Exp. Med. 2021, 21 (2), 167–179. https://doi.org/10.1007/s10238-020-00671-y.

(9) Gottlieb, R. L.; Nirula, A.; Chen, P.; Boscia, J.; Heller, B.; Morris, J.; Huhn, G.; Cardona, J.; Mocherla, B.; Stosor, V.; Shawa, I.; Kumar, P.; Adams, A. C.; Van Naarden, J.; Custer, K. L.; Durante, M.; Oakley, G.; Schade, A. E.; Holzer, T. R.; Ebert, P. J.; Higgs, R. E.; Kallewaard, N. L.; Sabo, J.; Patel, D. R.; Klekotka, P.; Shen, L.; Skovronsky, D. M. Effect of Bamlanivimab as Monotherapy or in Combination with Etesevimab on Viral Load in Patients with Mild to Moderate COVID-19: A Randomized Clinical Trial. JAMA - J. Am. Med. Assoc. 2021, 325 (7), 632–644. https://doi.org/10.1001/jama.2021.0202.

(10) Kokic, G.; Hillen, H. S.; Tegunov, D.; Dienemann, C.; Seitz, F.; Schmitzova, J.; Farnung, L.; Siewert, A.; Höbartner, C.; Cramer, P. Mechanism of SARS-CoV-2 Polymerase Stalling by Remdesivir. Nat. Commun. 2021, 12 (1), 1–7. https://doi.org/10.1038/s41467-020-20542-0.

(11) Elfiky, A. A. Ribavirin, Remdesivir, Sofosbuvir, Galidesivir, and Tenofovir against SARS-CoV-2 RNA Dependent RNA Polymerase (RdRp): A Molecular Docking Study. Life Sci. 2020, 253 (February). https://doi.org/10.1016/j.lfs.2020.117592.

(12) Beigel, J. H.; Tomashek, K. M.; Dodd, L. E.; Mehta, A. K.; Zingman, B. S.; Kalil, A. C.; Hohmann, E.; Chu, H. Y.; Luetkemeyer, A.; Kline, S.; Lopez de Castilla, D.; Finberg, R. W.; Dierberg, K.; Tapson, V.; Hsieh, L.; Patterson, T. F.; Paredes, R.; Sweeney, D. A.; Short, W. R.; Touloumi, G.; Lye, D. C.; Ohmagari, N.; Oh, M.; Ruiz-Palacios, G. M.; Benfield, T.; Fätkenheuer, G.; Kortepeter, M. G.; Atmar, R. L.; Creech, C. B.; Lundgren, J.; Babiker, A. G.; Pett, S.; Neaton, J. D.; Burgess, T. H.; Bonnett, T.; Green, M.; Makowski, M.; Osinusi, A.; Nayak, S.; Lane, H. C. Remdesivir for the Treatment of Covid-19 — Final Report. N. Engl. J. Med. 2020, 383 (19), 1813–1826. https://doi.org/10.1056/nejmoa2007764.

(13) Zhang, J.; Xie, B.; Hashimoto, K. Current Status of Potential Therapeutic Candidates for the COVID-19 Crisis. Brain. Behav. Immun. 2020. https://doi.org/https://doi.org/10.1016/j.bbi.2020.04.046.

(14) Mohammad Zadeh, N.; Mashinchi Asl, N. S.; Forouharnejad, K.; Ghadimi, K.; Parsa, S.; Mohammadi, S.; Omidi, A. Mechanism and Adverse Effects of COVID-19 Drugs: A Basic Review. Int. J. Physiol. Pathophysiol. Pharmacol. 2021, 13 (4), 102–109.

(15) Nguyen, N. T.; Chinn, J.; Nahmias, J.; Yuen, S.; Kirby, K. A.; Hohmann, S.; Amin, A. Outcomes and Mortality among Adults Hospitalized with COVID-19 at US Medical Centers. JAMA Netw. Open 2021, 4 (3), 20–23. https://doi.org/10.1001/jamanetworkopen.2021.0417.

(16) Naqvi, A. A. T.; Fatima, K.; Mohammad, T.; Fatima, U.; Singh, I. K.; Singh, A.; Atif, S. M.; Hariprasad, G.; Hasan, G. M.; Hassan, M. I. Insights into SARS-CoV-2 Genome, Structure, Evolution, Pathogenesis and Therapies: Structural Genomics Approach. BBA - Mol. Basis Dis. 2020, 1866, 165878. https://doi.org/https://doi.org/10.1016/j.bbadis.2020.165878.

(17) Hu, B.; Guo, H.; Zhou, P.; Shi, Z. L. Characteristics of SARS-CoV-2 and COVID-19. Nat. Rev. Microbiol. 2021, 19 (3), 141–154. https://doi.org/10.1038/s41579-020-00459-7.

(18) Mollica, V.; Rizzo, A.; Massari, F. The Pivotal Role of TMPRSS2 in Coronavirus Disease 2019 and Prostate Cancer. Futur. Oncol. 2020, 16 (27), 2029–2033. https://doi.org/10.2217/fon-2020-0571.

(19) Cohen, M. S. Monoclonal Antibodies to Disrupt Progression of Early Covid-19 Infection. N. Engl. J. Med. 2021, 384 (3), 289–291. https://doi.org/10.1056/nejme2033310.

(20) Fareh, M.; Zhao, W.; Hu, W.; Casan, J. M. L.; Kumar, A.; Symons, J.; Zerbato, J. M.; Fong, D.; Voskoboinik, I.; Ekert, P. G.; Rudraraju, R.; Purcell, D. F. J.; Lewin, S. R.; Trapani, J. A. Reprogrammed CRISPR-Cas13b Suppresses SARS-CoV-2 Replication and Circumvents Its Mutational Escape through Mismatch Tolerance. Nat. Commun. 2021, 12 (1). https://doi.org/10.1038/s41467-021-24577-9.

(21) Abbott, T. R.; Dhamdhere, G.; Liu, Y.; Lin, X.; Goudy, L.; Zeng, L.; Chemparathy, A.; Chmura, S.; Heaton, N. S.; Debs, R.; Pande, T.; Endy, D.; La Russa, M. F.; Lewis, D. B.; Qi, L. S. Development of CRISPR as an Antiviral Strategy to Combat SARS-CoV-2 and Influenza. Cell 2020, 181 (4), 865–876.e12. https://doi.org/10.1016/j.cell.2020.04.020.

(22) Idris, A.; Davis, A.; Supramaniam, A.; Acharya, D.; Kelly, G.; Tayyar, Y.; West, N.; Zhang, P.; McMillan, C. L. D.; Soemardy, C.; Ray, R.; O’Meally, D.; Scott, T. A.; McMillan, N. A. J.; Morris, K. V. A SARS-CoV-2 Targeted SiRNA-Nanoparticle Therapy for COVID-19. Mol. Ther. 2021, 29 (7), 2219–2226. https://doi.org/10.1016/j.ymthe.2021.05.004.

(23) Kanasty, R.; Dorkin, J. R.; Vegas, A.; Anderson, D. Delivery Materials for SiRNA Therapeutics. Nat. Mater. 2013, 12 (11), 967–977. https://doi.org/10.1038/nmat3765.

(24) Bost, J. P.; Barriga, H.; Holme, M. N.; Gallud, A.; Maugeri, M.; Gupta, D.; Lehto, T.; Valadi, H.; Esbjörner, E. K.; Stevens, M. M.; El-Andaloussi, S. Delivery of Oligonucleotide Therapeutics: Chemical Modifications, Lipid Nanoparticles, and Extracellular Vesicles. ACS Nano 2021, 15 (9), 13993–14021. https://doi.org/10.1021/acsnano.1c05099.

(25) Hegde, S.; Tang, Z.; Zhao, J.; Wang, J. Inhibition of SARS-CoV-2 by Targeting Conserved Viral RNA Structures and Sequences. Front. Chem. 2021, 9 (December), 1–11. https://doi.org/10.3389/fchem.2021.802766.

(26) Li, D. K.; Chung, R. T. Overview of Direct-Acting Antiviral Drugs and Drug Resistance of Hepatitis C Virus. In Hepatitis C Virus Protocols; Law, M., Ed.; Humana Press: New York, 2019; pp 3–32. https://doi.org/https://doi.org/10.1007/978-1-4939-8976-8_1.

(27) Pellestor, F.; Paulasova, P. The Peptide Nucleic Acids (PNAs), Powerful Tools for Molecular Genetics and Cytogenetics. Eur. J. Hum. Genet. 2004, 12 (9), 694–700. https://doi.org/10.1038/sj.ejhg.5201226.

(28) Li, C.; Callahan, A. J.; Phadke, K. S.; Bellaire, B.; Farquhar, C. E.; Zhang, G.; Schissel, C. K.; Mijalis, A. J.; Hartrampf, N.; Loas, A.; Verhoeven, D. E.; Pentelute, B. L. Automated Flow Synthesis of Peptide-PNA Conjugates. ACS Cent. Sci. 2021. https://doi.org/10.1021/acscentsci.1c01019.

(29) Upadhyay, A.; Ponzio, N. M.; Pandey, V. N. Immunological Response to Peptide Nucleic Acid and Its Peptide Conjugate Targeted to Transactivation Response (TAR) Region of HIV-1 RNA Genome. Oligonucleotides 2008, 18 (4), 329–335. https://doi.org/10.1089/oli.2008.0152.

(30) Habault, J.; Poyet, J. L. Recent Advances in Cell Penetrating Peptide-Based Anticancer Therapies. Molecules 2019, 24 (5), 1–17. https://doi.org/10.3390/molecules24050927.

(31) Meng, Z.; Lu, M. RNA Interference-Induced Innate Immunity, off-Target Effect, or Immune Adjuvant? Front. Immunol. 2017, 8 (MAR), 1–7. https://doi.org/10.3389/fimmu.2017.00331.

(32) Sioud, M. Induction of Inflammatory Cytokines and Interferon Responses by Double-Stranded and Single-Stranded SiRNAs Is Sequence-Dependent and Requires Endosomal Localization. J. Mol. Biol. 2005, 348 (5), 1079–1090. https://doi.org/10.1016/j.jmb.2005.03.013.

(33) Tan, A.; Hong, L.; Du, J. D.; Boyd, B. J. Self-Assembled Nanostructured Lipid Systems: Is There a Link between Structure and Cytotoxicity? Adv. Sci. 2019, 6 (3). https://doi.org/10.1002/advs.201801223.

(34) Gretebeck, L. M.; Subbarao, K. Animal Models for SARS and MERS Coronaviruses. Curr. Opin. Virol. 2015, 13, 123–129. https://doi.org/http://dx.doi.org/10.1016/j.coviro.2015.06.009.

(35) Muñoz-Fontela, C.; Dowling, W. E.; Funnell, S. G. P.; Gsell, P. S.; Riveros-Balta, A. X.; Albrecht, R. A.; Andersen, H.; Baric, R. S.; Carroll, M. W.; Cavaleri, M.; Qin, C.; Crozier, I.; Dallmeier, K.; de Waal, L.; de Wit, E.; Delang, L.; Dohm, E.; Duprex, W. P.; Falzarano, D.; Finch, C. L.; Frieman, M. B.; Graham, B. S.; Gralinski, L. E.; Guilfoyle, K.; Haagmans, B. L.; Hamilton, G. A.; Hartman, A. L.; Herfst, S.; Kaptein, S. J. F.; Klimstra, W. B.; Knezevic, I.; Krause, P. R.; Kuhn, J. H.; Le Grand, R.; Lewis, M. G.; Liu, W. C.; Maisonnasse, P.; McElroy, A. K.; Munster, V.; Oreshkova, N.; Rasmussen, A. L.; Rocha-Pereira, J.; Rockx, B.; Rodríguez, E.; Rogers, T. F.; Salguero, F. J.; Schotsaert, M.; Stittelaar, K. J.; Thibaut, H. J.; Tseng, C. Te; Vergara-Alert, J.; Beer, M.; Brasel, T.; Chan, J. F. W.; García-Sastre, A.; Neyts, J.; Perlman, S.; Reed, D. S.; Richt, J. A.; Roy, C. J.; Segalés, J.; Vasan, S. S.; Henao-Restrepo, A. M.; Barouch, D. H. Animal Models for COVID-19. Nature 2020, 586 (7830), 509–515. https://doi.org/10.1038/s41586-020-2787-6.

(36) Wang, Q. hACE2 Transgenic Mouse Model for Coronavirus (COVID-19) Research https://www.jax.org/news-and-insights/2020/february/introducing-mouse-model-for-corona-virus# (accessed Apr 13, 2020).

(37) Pires, A.; Fortuna, A.; Alves, G.; Falcão, A. Intranasal Drug Delivery: How, Why and What For? J. Pharm. Pharm. Sci. 2009, 12 (3), 288–311. https://doi.org/10.18433/j3nc79.

(38) Al Shoyaib, A.; Archie, S. R.; Karamyan, V. T. Intraperitoneal Route of Drug Administration: Should It Be Used in Experimental Animal Studies? Pharm. Res. 2020, 37 (1), 1–30. https://doi.org/10.1007/s11095-019-2745-x.

(39) Grossoehme, N. E.; Li, L.; Keane, S. C.; Liu, P.; Dann, C. E.; Leibowitz, J. L.; Giedroc, D. P. Coronavirus N Protein N-Terminal Domain (NTD) Specifically Binds the Transcriptional Regulatory Sequence (TRS) and Melts TRS-CTRS RNA Duplexes. J. Mol. Biol. 2009, 394 (3), 544–557. https://doi.org/10.1016/j.jmb.2009.09.040.

(40) Good, L.; Awasthi, S. K.; Dryselius, R.; Larsson, O.; Nielsen, P. E. Bactericidal Antisense Effects of Peptide - PNA Conjugates. Nat. Biotechnol. 2001, 19 (4), 360–364. https://doi.org/10.1038/86753.

(41) Kilså Jensen, K.; Ørum, H.; Nielsen, P. E.; Nordén, B. Kinetics for Hybridization of Peptide Nucleic Acids (PNA) with DNA and RNA Studied with the BIAcore Technique. Biochemistry 1997, 36 (16), 5072–5077. https://doi.org/10.1021/bi9627525.

(42) Zaias, J.; Mineau, M.; Cray, C.; Yoon, D.; Altman, N. H. Reference Values for Serum Proteins of Common Laboratory Rodent Strains. J. Am. Assoc. Lab. Anim. Sci. 2009, 48 (4), 387–390.

(43) Natsume, M.; Tsuji, H.; Harada, A.; Akiyama, M.; Yano, T.; Ishikura, H.; Nakanishi, I.; Matsushima, K.; Kaneko, S. I.; Mukaida, N. Attenuated Liver Fibrosis and Depressed Serum Albumin Levels in Carbon Tetrachloride-Treated IL-6-Deficient Mice. J. Leukoc. Biol. 1999, 66 (4), 601–608. https://doi.org/10.1002/jlb.66.4.601.

(44) The Jackson Laboratory. Mouse Phenome Database: BALB/CByJ.

(45) Rosseels, V.; Nazé, F.; De Craeye, S.; Francart, A.; Kalai, M.; Van Gucht, S. A Non-Invasive Intranasal Inoculation Technique Using Isoflurane Anesthesia to Infect the Brain of Mice with Rabies Virus. J. Virol. Methods 2011, 173 (1), 127–136. https://doi.org/10.1016/j.jviromet.2011.01.019.

(46) Rao, G. V. S.; Tinkle, S.; Weissman, D. N.; Antonini, J. M.; Kashon, M. L.; Salmen, R.; Battelli, L. A.; Willard, P. A.; Hubbs, A. F.; Hoover, M. D. Efficacy of a Technique for Exposing the Mouse Lung to Particles Aspirated from the Pharynx. J. Toxicol. Environ. Heal. - Part A 2003, 66 (15), 1441–1452. https://doi.org/10.1080/15287390306417.

(47) Geary, R. S.; Norris, D.; Yu, R.; Bennett, C. F. Pharmacokinetics, Biodistribution and Cell Uptake of Antisense Oligonucleotides. Advanced Drug Delivery Reviews. 2015. https://doi.org/10.1016/j.addr.2015.01.008.

(48) Smith, R. A.; Miller, T. M.; Yamanaka, K.; Monia, B. P.; Condon, T. P.; Hung, G.; Lobsiger, C. S.; Ward, C. M.; McAlonis-Downes, M.; Wei, H.; Wancewicz, E. V.; Bennett, C. F.; Cleveland, D. W. Antisense Oligonucleotide Therapy for Neurodegenerative Disease. J. Clin. Invest. 2006, 116 (8), 2290–2296. https://doi.org/10.1172/JCI25424.

(49) Yu, R. Z.; Lemonidis, K. M.; Graham, M. J.; Matson, J. E.; Crooke, R. M.; Tribble, D. L.; Wedel, M. K.; Levin, A. A.; Geary, R. S. Cross-Species Comparison of in Vivo PK/PD Relationships for Second-Generation Antisense Oligonucleotides Targeting Apolipoprotein B-100. Biochem. Pharmacol. 2009, 77 (5), 910–919. https://doi.org/10.1016/j.bcp.2008.11.005.

(50) Altmann, K.-H.; Dean, N. M.; Fabbro, D.; Freier, S. M.; Geiger, T.; Haner, R.; Husken, D.; Martin, P.; Monia, B. P.; Muller, M.; Natt, F.; Nicklin, P.; Phillips, J.; Pieles, U.; Sasmor, H.; Moser, H. E. Second Generation of Antisense Oligonucleotides: From Nuclease Resistance to Biological Efficacy in Animals. Chimia (Aarau). 1996, 50 (4), 168–176. https://doi.org/https://doi.org/10.2533/chimia.1996.168.

(51) Geary, R. S.; Wancewicz, E.; Matson, J.; Pearce, M.; Siwkowski, A.; Swayze, E.; Bennett, F. Effect of Dose and Plasma Concentration on Liver Uptake and Pharmacologic Activity of a 2’-Methoxyethyl Modified Chimeric Antisense Oligonucleotide Targeting PTEN. Biochem. Pharmacol. 2009, 78 (3), 284–291. https://doi.org/10.1016/j.bcp.2009.04.013.

(52) Kaur, N.; Aditya, R. N.; Singh, A.; Kuo, T. R. Biomedical Applications for Gold Nanoclusters: Recent Developments and Future Perspectives. Nanoscale Res. Lett. 2018, 13. https://doi.org/10.1186/s11671-018-2725-9.

(53) Tian, L.; Qiang, T.; Liang, C.; Ren, X.; Jia, M.; Zhang, J.; Li, J.; Wan, M.; YuWen, X.; Li, H.; Cao, W.; Liu, H. RNA-Dependent RNA Polymerase (RdRp) Inhibitors: The Current Landscape and Repurposing for the COVID-19 Pandemic. Eur. J. Med. Chem. 2021, 213, 113201. https://doi.org/10.1016/j.ejmech.2021.113201.

(54) World Health Organization. Antiviral Drugs That Are Approved or Under Evaluation for the Treatment of COVID-19. COVID-19 Treat. Guidel. 2020, 19, 47–88.

(55) McDonald, J. T.; Enguita, F. J.; Taylor, D.; Griffin, R. J.; Priebe, W.; Emmett, M. R.; Sajadi, M. M.; Harris, A. D.; Clement, J.; Dybas, J. M.; Aykin-Burns, N.; Guarnieri, J. W.; Singh, L. N.; Grabham, P.; Baylin, S. B.; Yousey, A.; Pearson, A. N.; Corry, P. M.; Saravia-Butler, A.; Aunins, T. R.; Sharma, S.; Nagpal, P.; Meydan, C.; Foox, J.; Mozsary, C.; Cerqueira, B.; Zaksas, V.; Singh, U.; Wurtele, E. S.; Costes, S. V.; Davanzo, G. G.; Galwano, D.; Paccanaro, A.; Meinig, S. L.; Hagan, R. S.; Bowman, N. M.; UNC COVID-19 Pathobiology Consortium; Wolfgang, M. C.; Altinok, S.; Sapoval, N.; Treangen, T. J.; Moraes-Vieira, P. M.; Vanderburg, C.; Wallace, D. C.; Schisler, J. C.; Mason, C. E.; Chatterjee, A.; Meller, R.; Beheshti, A. Role of MiR-2392 in Driving SARS-CoV-2 Infection. Cell Rep. 2021, 37 (October), 1–17. https://doi.org/https://doi.org/10.1016/j.celrep.2021.109839.

(56) Eller, K. A.; Aunins, T. R.; Courtney, C. M.; Campos, J. K.; Otoupal, P. B.; Erickson, K. E.; Madinger, N. E.; Chatterjee, A. Facile Accelerated Specific Therapeutic (FAST) Platform Develops Antisense Therapies to Counter Multidrug-Resistant Bacteria. Commun. Biol. 2021, 4 (331), 1–13. https://doi.org/10.1038/s42003-021-01856-1.

(57) Aunins, T. R.; Erickson, K. E.; Chatterjee, A. Transcriptome-Based Design of Antisense Inhibitors Potentiates Carbapenem Efficacy in CRE Escherichia Coli. Proc. Natl. Acad. Sci. U. S. A. 2020, 117 (48), 30699–30709. https://doi.org/10.1073/pnas.1922187117.

(58) Eldrup, A. B.; Nielsen, P. E. Peptide Nucleic Acids; 1999. https://doi.org/10.1016/s1874-5113(99)80009-4.

(59) Daniloski, Z.; Jordan, T. X.; Wessels, H.-H.; Hoagland, D. A.; Kasela, S.; Legut, M.; Maniatis, S.; Mimitou, E. P.; Lu, L.; Geller, E.; Danziger, O.; Rosenberg, B. R.; Phatnani, H.; Smibert, P.; Lappalainen, T.; TenOever, B. R.; Sanjana, N. E. Identification of Required Host Factors for SARS-CoV-2 Infection in Human Cells. Cell 2021, 184 (January), 92–105. https://doi.org/https://doi.org/10.1016/j.cell.2020.10.030.

(60) Cardillo, G. Four Parameters Logistic Regression - There and Back Again. Mathworks File Exchange 2012.

(61) Thompson, J. D.; Kornbrust, D. J.; Foy, J. W. D.; Solano, E. C. R.; Schneider, D. J.; Feinstein, E.; Molitoris, B. A.; Erlich, S. Toxicological and Pharmacokinetic Properties of Chemically Modified SiRNAs Targeting P53 RNA Following Intravenous Administration. Nucleic Acid Ther. 2012, 22 (4), 255–264. https://doi.org/10.1089/nat.2012.0371.

(62) Gao, S.; Dagnaes-Hansen, F.; Nielsen, E. J. B.; Wengel, J.; Besenbacher, F.; Howard, K. A.; Kjems, J. The Effect of Chemical Modification and Nanoparticle Formulation on Stability and Biodistribution of SiRNA in Mice. Mol. Ther. 2009, 17 (7), 1225–1233. https://doi.org/10.1038/mt.2009.91.

(63) Hasgall, P.; Di Gennaro, F.; Baumgartner, C.; Neufeld, E.; Lloyd, B.; Gosselin, M.; Payne, D.; Klingenböck, A.; Kuster, N. IT’IS Database for thermal and electromagnetic parameters of biological tissues https://itis.swiss/virtual-population/tissue-properties/. https://doi.org/10.13099/VIP21000-04-0.

