## Supporting information for "Safety and biodistribution of Nanoligomers^™^ targeting SARS-CoV-2 genome for treatment of COVID-19"

### Contents


| Supplemental Tables | Page |
|---|---|
| Table S1 | S3 |
| Table S2 | S4 |
| **Supplemental Figures** | **Page** |
| Figure S1 | S5 |
| Figure S2 | S6 |
| Figure S3 | S7 |
| Figure S4 | S8 |
| Figure S5 | S9 |
| Figure S6 | S10 |
| Figure S7 | S11 |
| Figure S8 | S12 |
| Figure S9 | S13 |
| Figure S10 | S14 |
| Figure S11 | S15 |

**Table S1:**

| Administration | Time | Dose | TNF-α | IL-6 |
|---|---|---|---|---|
| Intranasal | 5 days | 1 mg/kg | Below level of detection | Below level of detection |
| Intranasal | 5 days | 2 mg/kg | Below level of detection | Below level of detection |
| Intranasal | 5 days | 5 mg/kg | Below level of detection | Below level of detection |
| Intranasal | 5 days | 10 mg/kg | Below level of detection | Below level of detection |
| Intraperitoneal | 5 days | 10 mg/kg | Below level of detection | Below level of detection |
| Intravenous | 5 days | 1 mg/kg | Below level of detection | Below level of detection |
| Intravenous | 5 days | 2 mg/kg | Below level of detection | Below level of detection |
| Intravenous | 5 days | 5 mg/kg | Below level of detection | Below level of detection |
| Intravenous | 5 days | 10 mg/kg | Below level of detection | Below level of detection |

**Table S1:** Serum from mice treated with SBCoV202 showed no detectable levels of TNF-α or IL-6 in ELISAs five days later, regardless of dose and administration route.

**Table S2:**

| Administration | Time | Dose | TNF-α | IL-6 |
|---|---|---|---|---|
| Intranasal | 1 hour | 10 mg/kg | Below level of detection | Below level of detection |
| Intranasal | 3 hours | 10 mg/kg | Below level of detection | Below level of detection |
| Intranasal | 6 hours | 10 mg/kg | Below level of detection | Below level of detection |
| Intranasal | 24 hours | 10 mg/kg | Below level of detection | Below level of detection |
| Intraperitoneal | 1 hour | 10 mg/kg | Below level of detection | Below level of detection |
| Intraperitoneal | 3 hours | 10 mg/kg | Below level of detection | Below level of detection |
| Intraperitoneal | 6 hours | 10 mg/kg | Below level of detection | Below level of detection |
| Intraperitoneal | 24 hours | 10 mg/kg | Below level of detection | Below level of detection |
| Intravenous | 1 hour | 10 mg/kg | Below level of detection | Below level of detection |
| Intravenous | 3 hours | 10 mg/kg | Below level of detection | Below level of detection |
| Intravenous | 6 hours | 10 mg/kg | Below level of detection | Below level of detection |
| Intravenous | 24 hours | 10 mg/kg | Below level of detection | Below level of detection |

**Table S2:** Serum from mice treated with SBCoV202 showed no detectable levels of TNF-α or IL-6 in ELISAs within 24 hours, regardless of administration route.

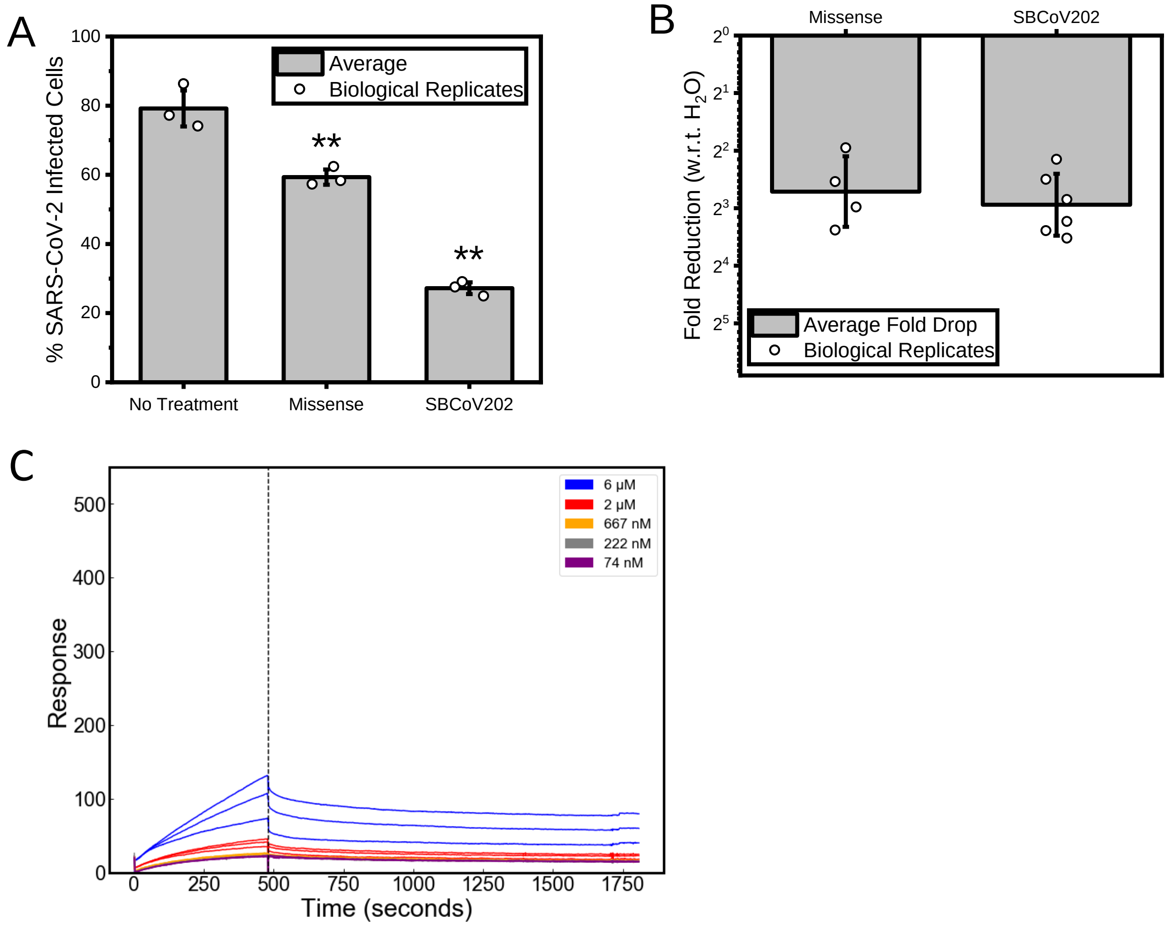


**Figure S1:** (**A**) After treatment with 10 μM SBCoV202, human A549-hACE2 lung epithelial cells infected with SARS-CoV-2, then fixed and immunostained for viral nucleocapsid protein, showed significant decrease in percent infection compared to untreated cells. Cells treated with 10 μM missense Nanoligomer showed some reduction of infection, but reduction was most prevalent with cells treated with SBCoV202. (**B**) Quantitative reverse-transcription polymerase chain reaction (qRT-PCR) showed expression of viral mRNA was reduced in infected cells treated with 10 μM Nanoligomers compared to a water control. Expression reduction was greater in cells treated with SBCoV202 than missense Nanoligomers. (**D**) Surface plasmon resonance (SPR) analysis shows that the missense control Nanoligomer does not bind the target, based on the low magnitude of response. P-value indicated by * < 0.05, ** < 0.01, *** < 0.001, otherwise p > 0.05.

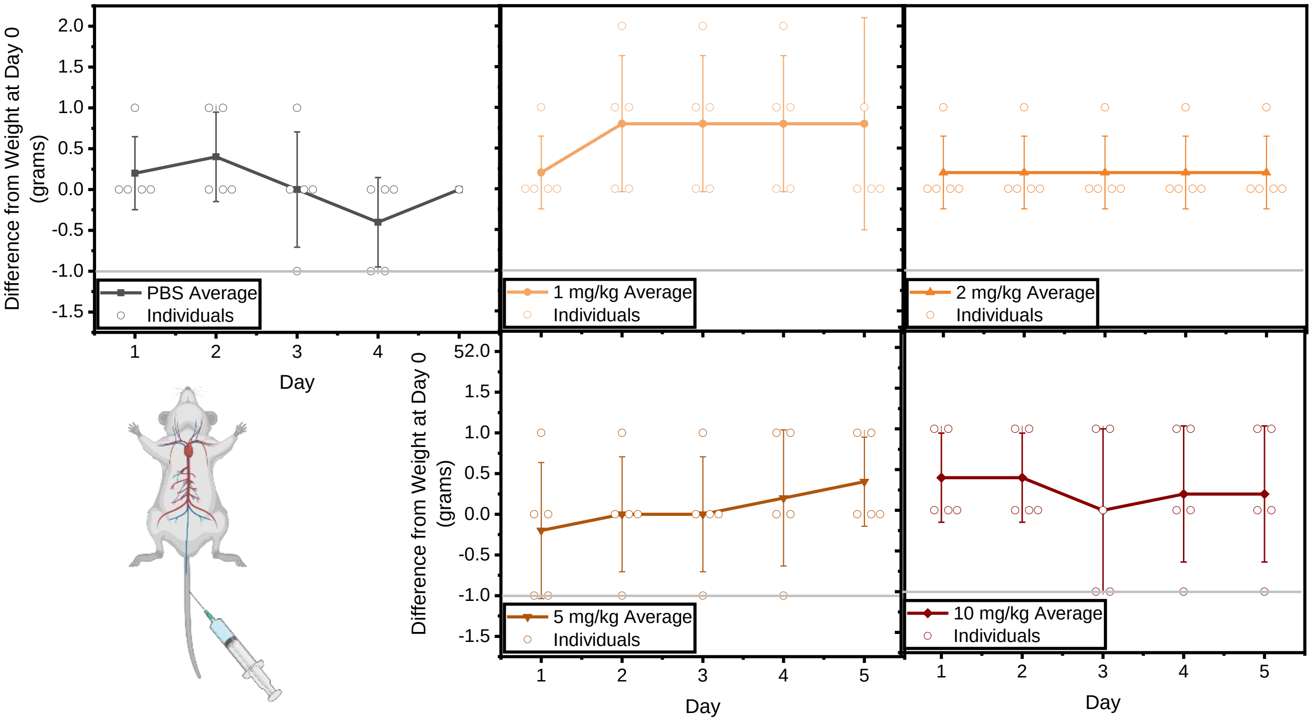


**Figure S2:** Intravenous administration of SBCoV202 did not cause weight loss in mice. Mice were administered with an injection of SBCoV202 into the lateral tail vein (visualized in bottom left) and monitored for 5 days. Weight did not drop more than 1 gram below starting weight for any mouse. The horizontal gray line is used to indicate 1 gram below the weight on Day 0 of the study. The p value was greater than 0.05 for treated mice at all timepoints compared to the PBS control group.

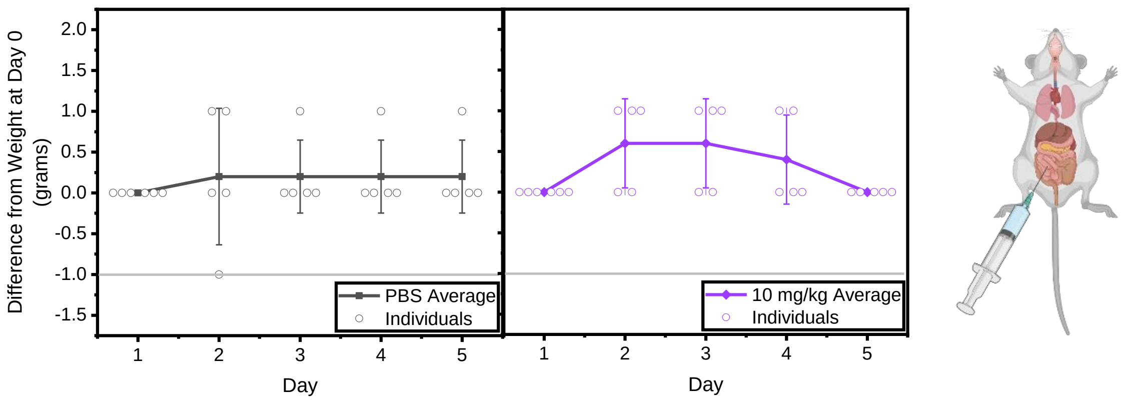


**Figure S3:** Intraperitoneal administration of SBCoV202 did not cause weight loss in mice. Mice were administered with an injection of SBCoV202 into the peritoneal space (visualized on right) and monitored for 5 days. Weight did not drop more than 1 gram below starting weight for any mouse. The horizontal gray line is used to indicate 1 gram below the weight on Day 0 of the study. The p value was greater than 0.05 for treated mice at all timepoints compared to the PBS control group.

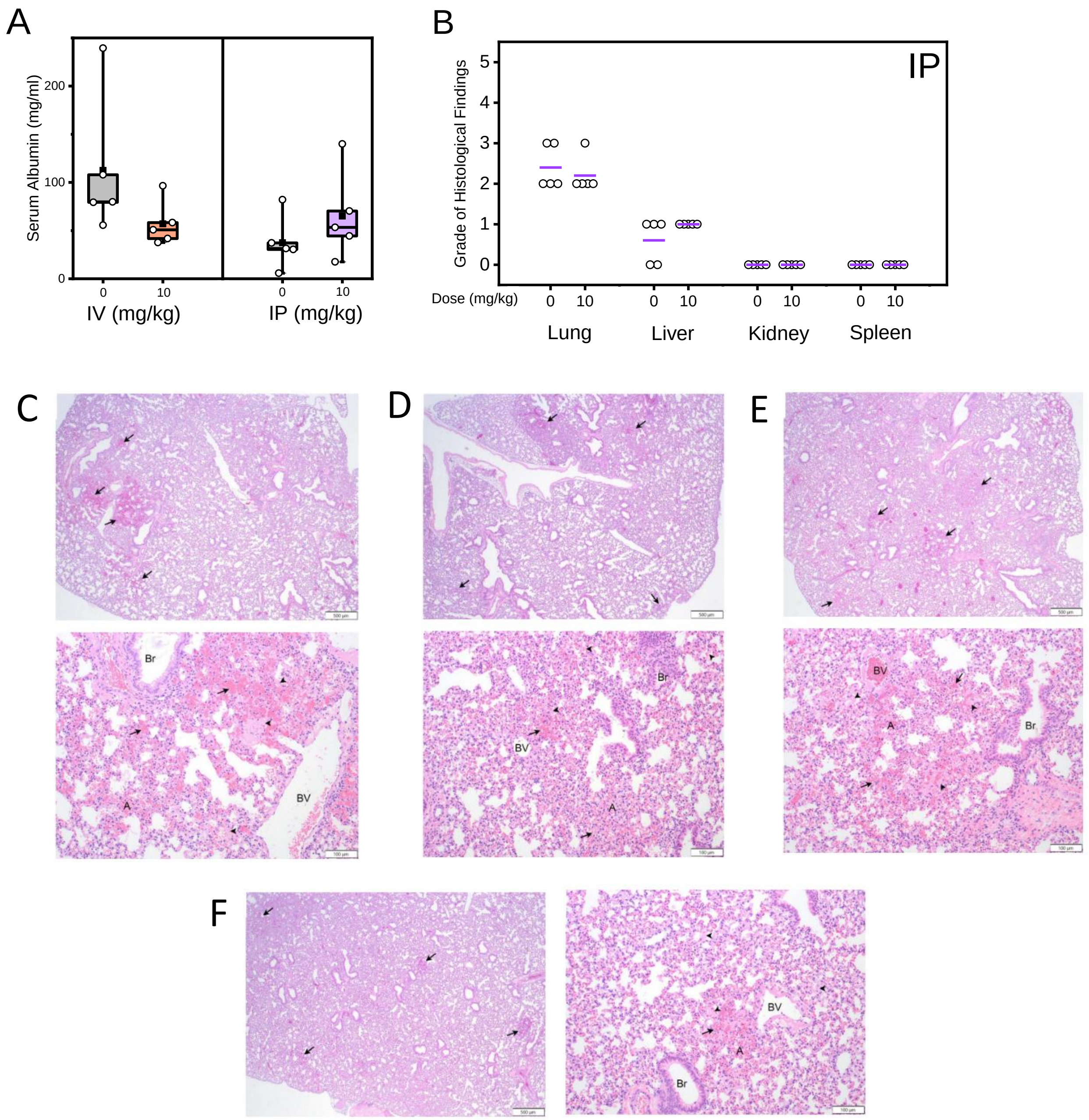


**Figure S4:** Administration of SBCoV202 did not result in significant changes to albumin levels in serum (**A**) or morphology of lung, liver, kidney, or spleen based on histology with H&E staining (**B**), in intraperitoneally injected mice. Histology was graded on a 0-5 scale with 0=absent, 1=minimal, 2=mild, 3=moderate, 4=marked, 5=severe. Colored bars represent mean. $p > 0.05$ for all points. Multifocal regions of hemorrhage (arrows) are observed throughout the parenchyma in lungs of mice treated with IV PBS (**C**), IV SBCoV202 (**D**), IP PBS (**E**), or IP SBCoV202 (**F**), often adjacent to blood vessels (BV), and characterized by free red blood cells within alveoli (A) and bronchioles (Br). Hemorrhage is accompanied by eosinophilic proteinaceous material (fibrin; arrowheads) that expands alveolar septa and fills some alveolar spaces. Upper images in **C**-**E** and left image in **F** are taken at 20x; lower images in **C**-**E** and right image in **F** are taken at 100x.

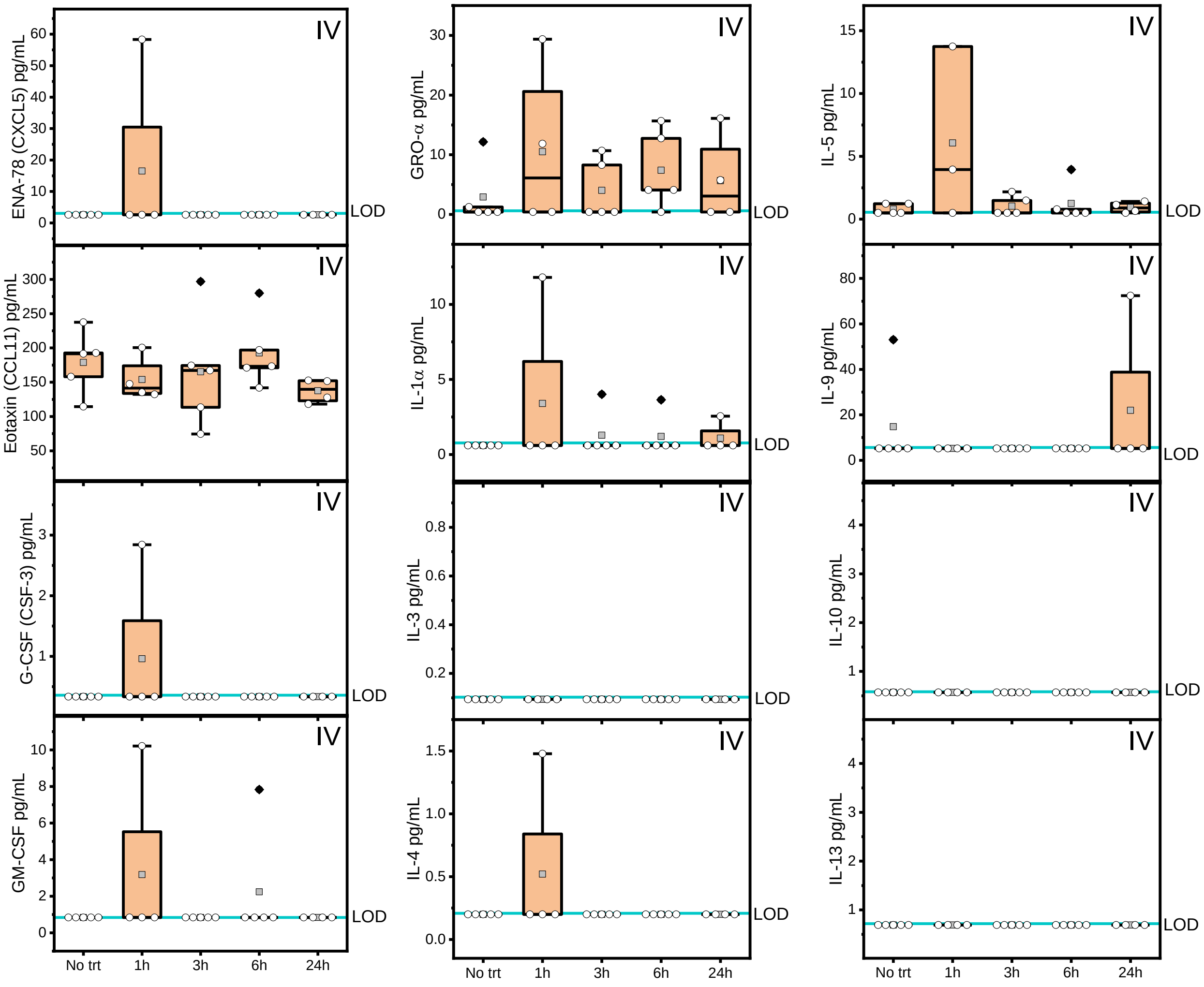


**Figure S5:** Measurements from a 36-cytokine/chemokine panel performed on serum samples from mice treated with SBCoV202 via intravenous (IV) injection showed cytokines largely below the limit of detection (LOD) of the assay, namely ENA-78, G-CSF, GM-CSF, IL-1α, IL-3, IL-4, IL-9, IL-10, and IL-13. The LOD is represented with a blue line. Levels of eotaxin, GRO-α, and IL-5 were comparable to the PBS control group. All groups showed $p > 0.05$ compared to the PBS control group.

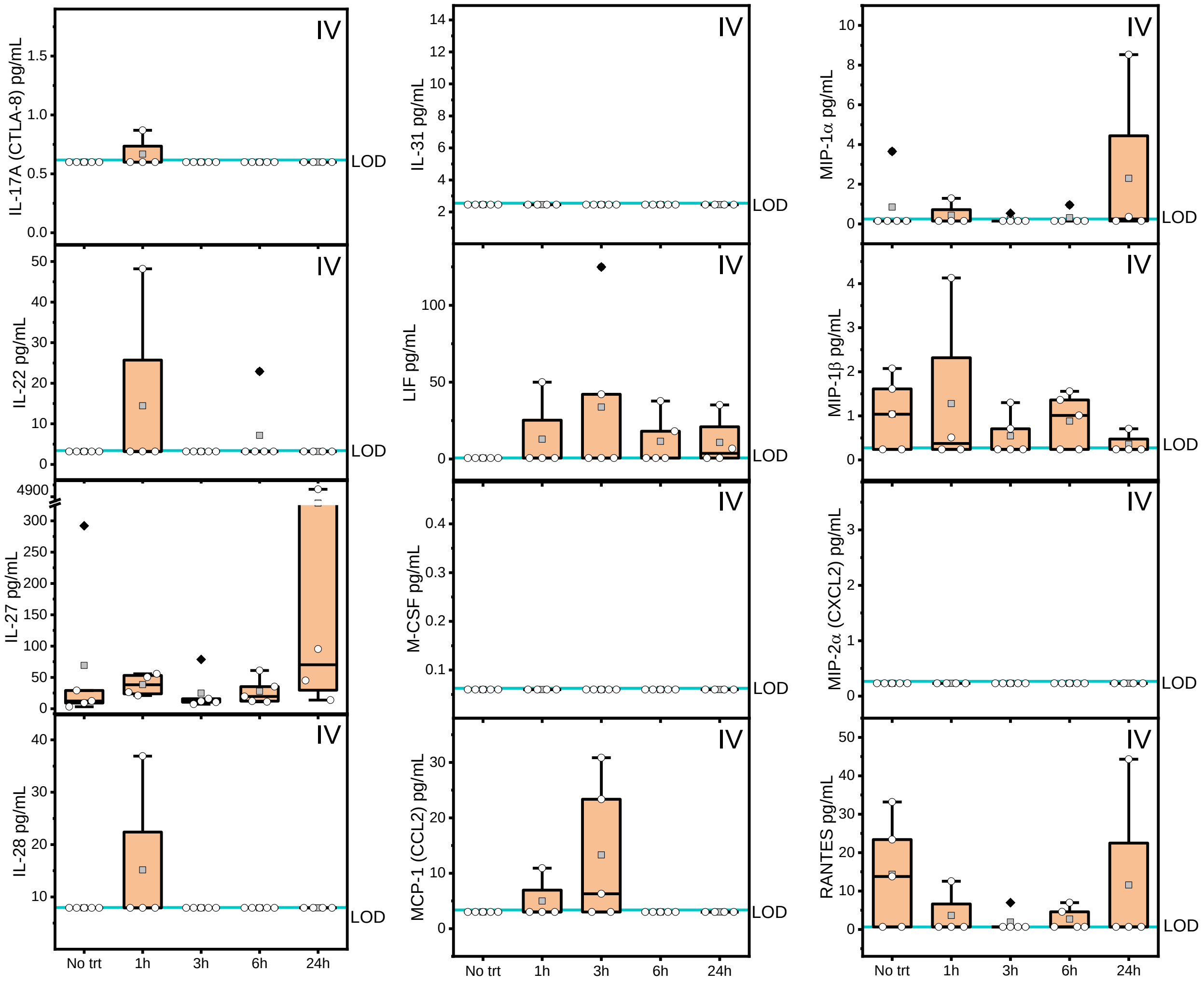


**Figure S6:** Measurements from a 36-cytokine/chemokine panel performed on serum samples from mice treated with SBCoV202 via intravenous (IV) injection showed cytokines largely below the limit of detection (LOD) of the assay, namely IL-17A, IL-22, IL-28, IL-31, M-CSF, and MIP-2α. The LOD is represented with a blue line. Levels of IL-27, LIF, MCP-1, MIP-1α, MIP-1β, and RANTES were comparable to the PBS control group. All groups showed $p > 0.05$ compared to the PBS control group.

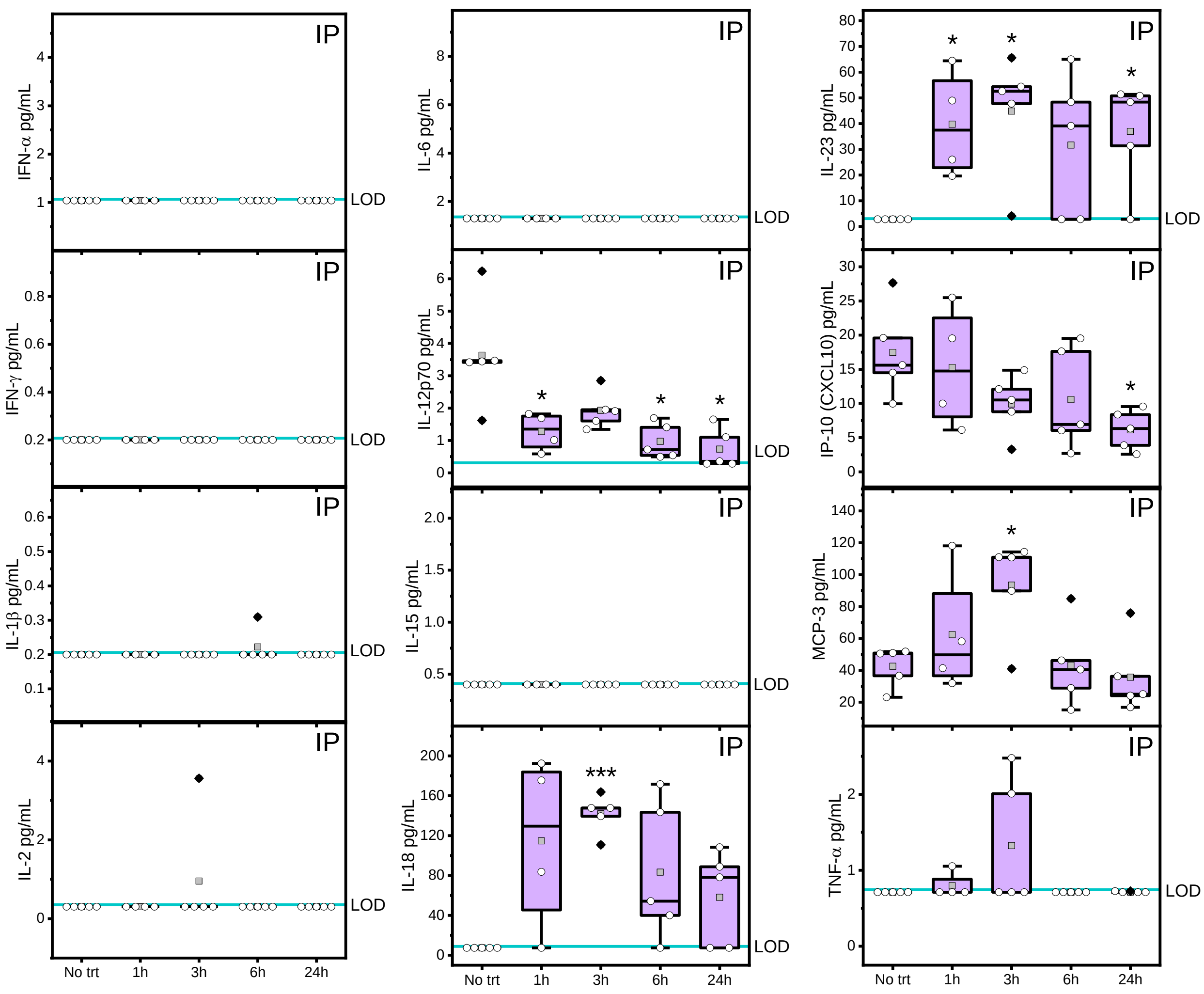


**Figure S7:** Key measurements from a 36-cytokine/chemokine panel performed on serum samples from mice treated with SBCoV202 via intraperitoneal (IP) injection showed key cytokines largely below the limit of detection (LOD) of the assay, namely IFN-α, IFN-γ, IL-1β, IL-2, IL-6, IL-15, and TNF-α. The LOD is represented with a blue line. Levels of IL-12p70 and IP-10 were lower than the PBS control group at some timepoints. Only IL-18, IL-23, and MCP-3 levels were significantly increased compared to the control. * indicates $p < 0.05$, *** indicates $p < 0.001$. All other groups showed $p > 0.05$ compared to the PBS control group.

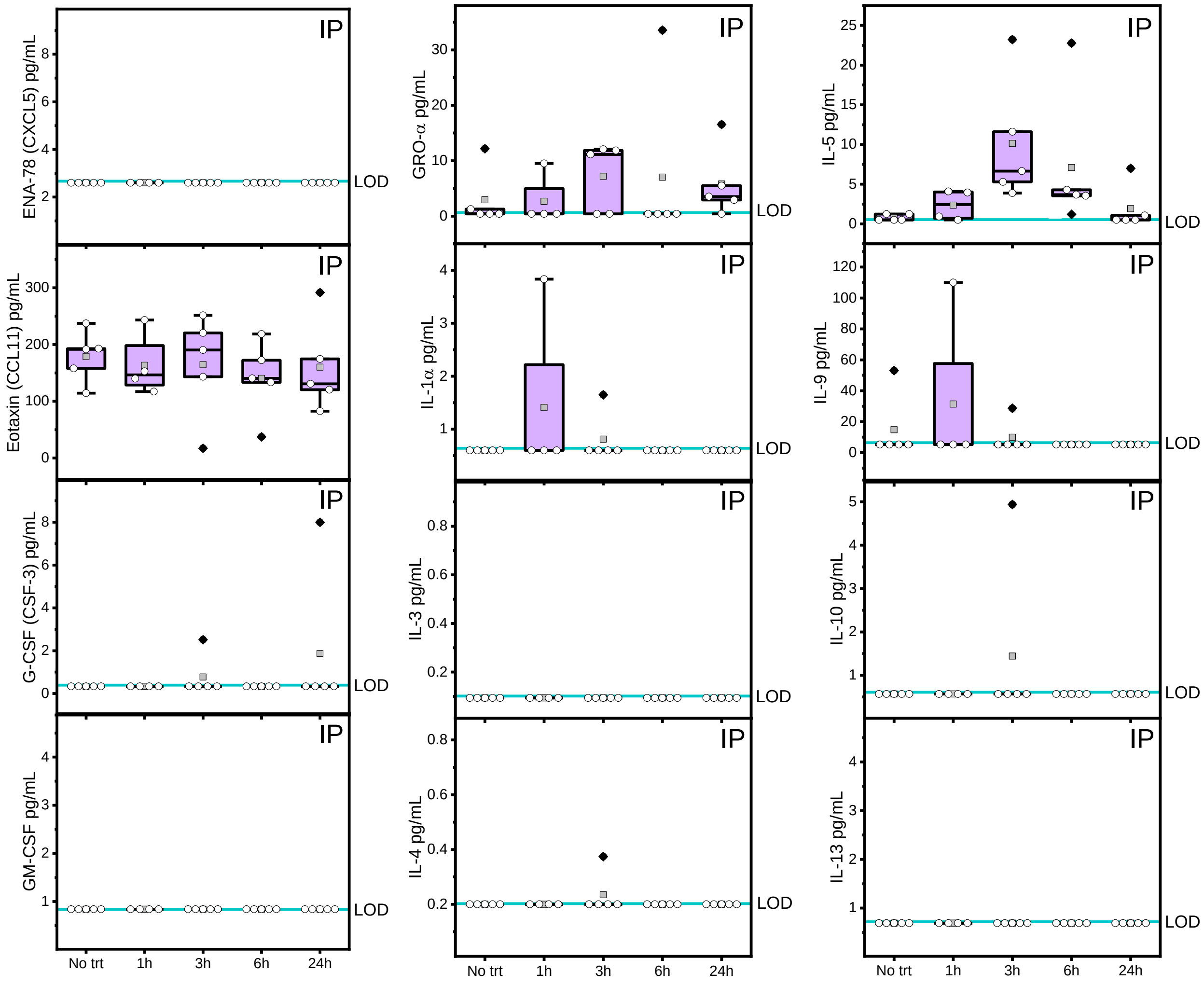


**Figure S8:** Measurements from a 36-cytokine/chemokine panel performed on serum samples from mice treated with SBCoV202 via intraperitoneal (IP) injection showed cytokines largely below the limit of detection (LOD) of the assay, namely ENA-78, G-CSF, GM-CSF, IL-1α, IL-3, IL-4, IL-9, IL-10, and IL-13. The LOD is represented with a blue line. Levels of eotaxin, GRO-α, and IL-5 were comparable to the PBS control group. All groups showed $p > 0.05$ compared to the PBS control group.

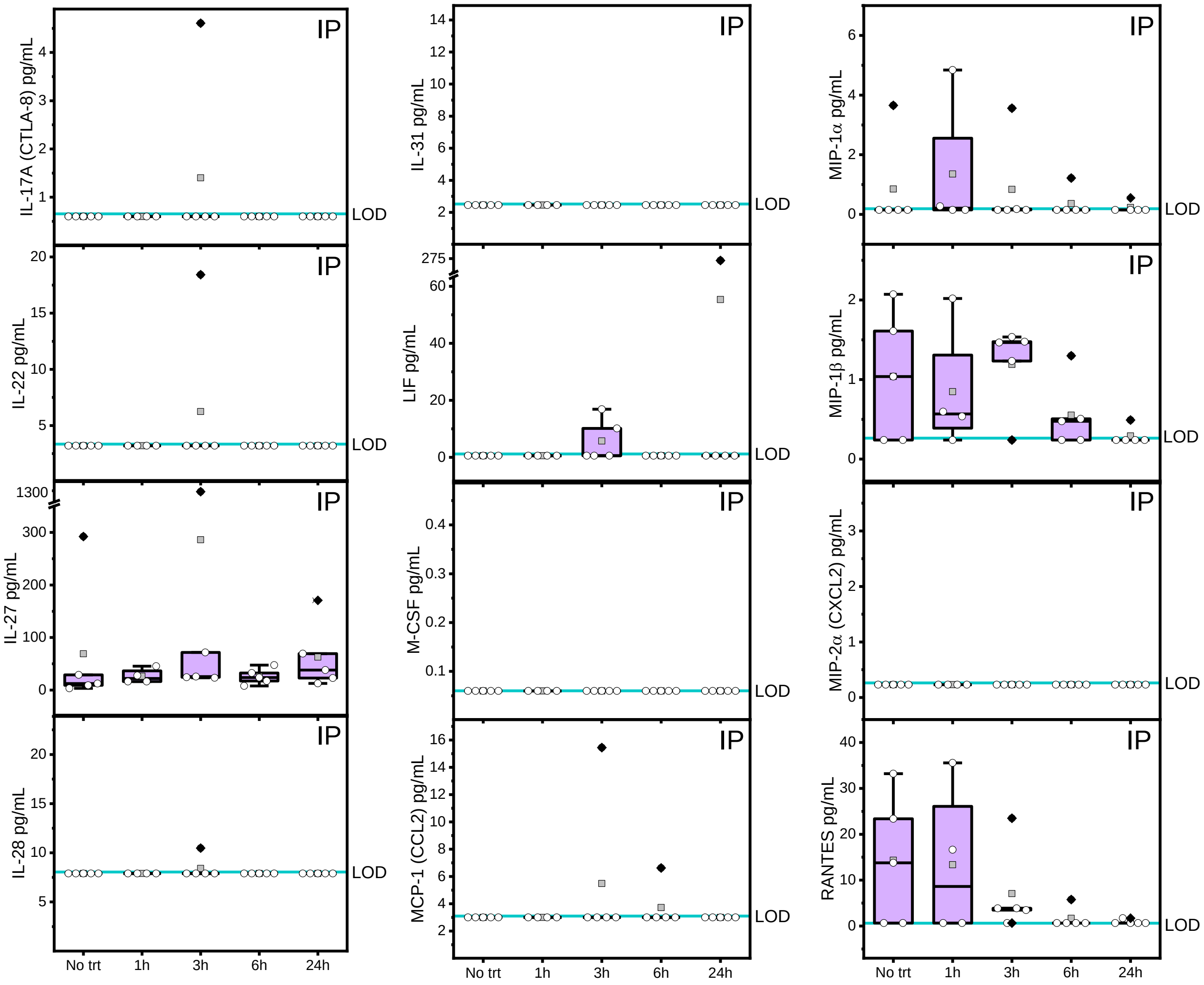


**Figure S9:** Measurements from a 36-cytokine/chemokine panel performed on serum samples from mice treated with SBCoV202 via intraperitoneal (IP) injection showed cytokines largely below the limit of detection (LOD) of the assay, namely IL-17A, IL-22, IL-28, IL-31, LIF, M-CSF, MCP-1, and MIP-2α. The LOD is represented with a blue line. Levels of IL-27, MIP-1α, MIP-1β, and RANTES were comparable to the PBS control group. All groups showed $p > 0.05$ compared to the PBS control group.

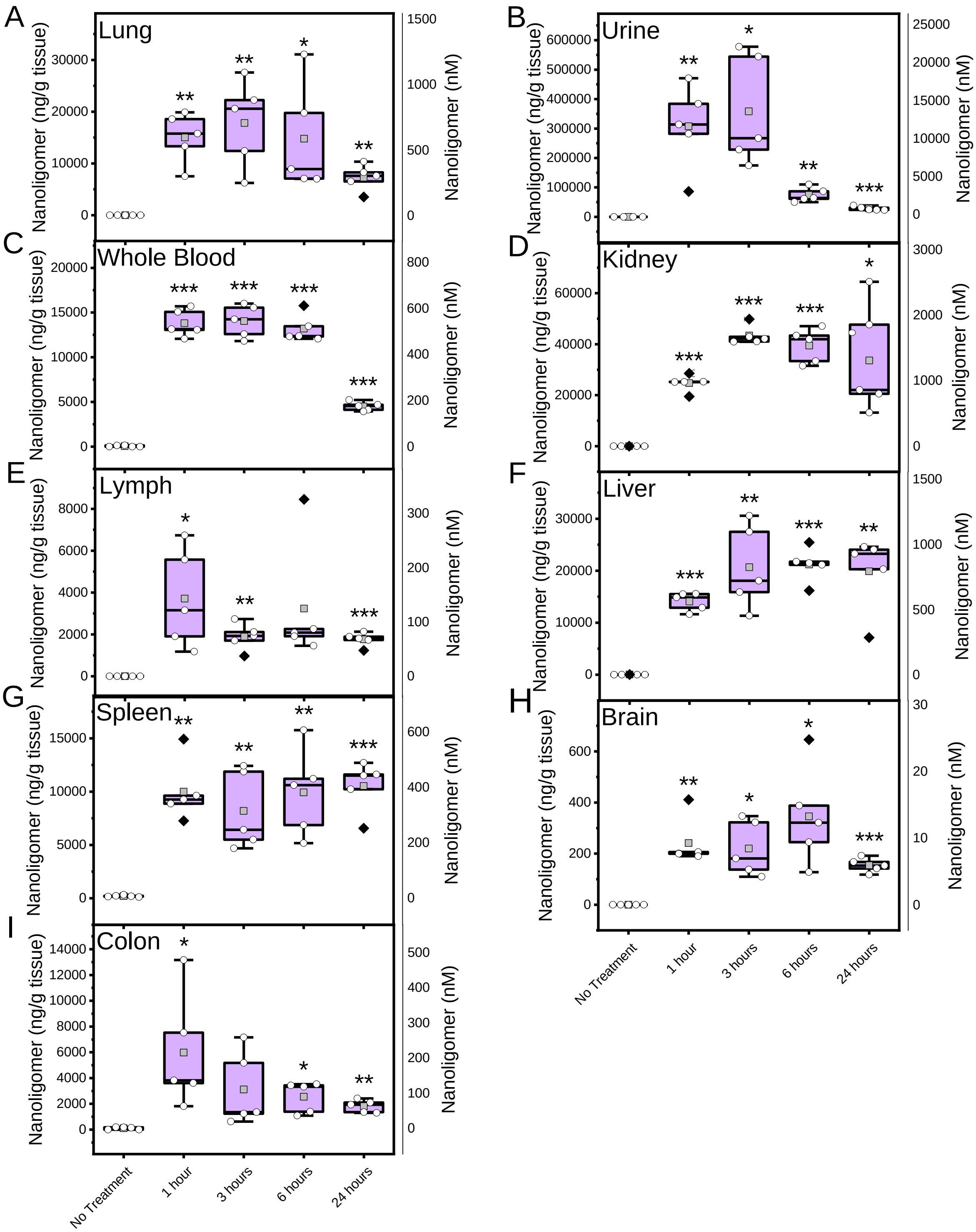


**Figure S10:** SBCoV202 biodistribution after intraperitoneal administration showed SBCoV202 (shown as nanograms per gram of tissue on left, nanomolar on right) reaching the lungs (**A**), being cleared by the renal system (**D**), and excreted by the urinary system (**B**). SBCoV202 was circulated in the whole blood (**C**) and detected in lymph (**E**) and liver (**F**) tissue. Levels in the spleen (**G**), brain (**H**), and colon (**I**) were lower than other organs. P-value indicated by * < 0.05, ** < 0.01, *** < 0.001, otherwise p > 0.05.

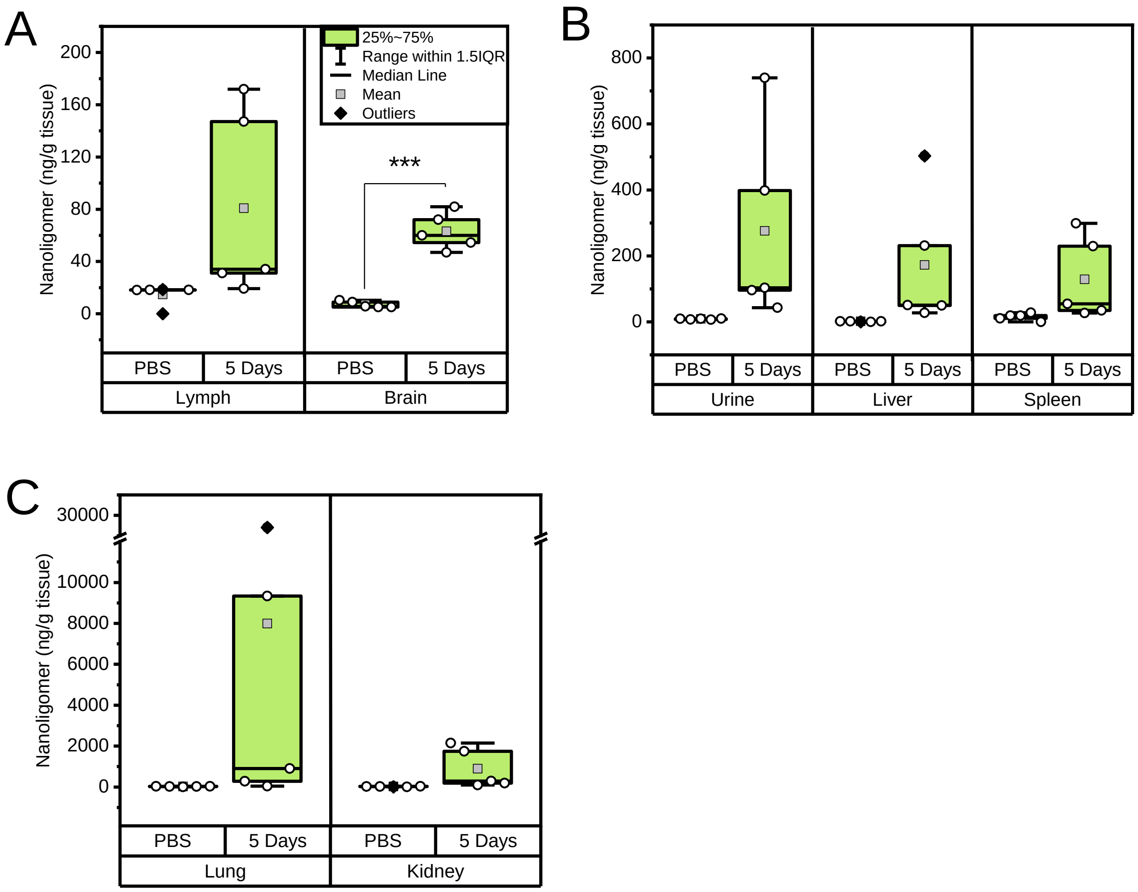


**Figure S11:** SBCoV202 biodistribution five days after 10 mg/kg intranasal administration showed SBCoV202 are largely no longer present in the lymph or brain tissue (**A**), present to a moderate degree only in some mice in urine, liver, and spleen (**B**), and still present at higher concentrations in only two mice in lung (**C**). P-value indicated by *** < 0.001, otherwise p > 0.05.
